# The long noncoding RNA *Hand2as* orchestrates heart development through regulation of precise expression of *HAND2*

**DOI:** 10.1101/475723

**Authors:** Xue Han, Jiejie Zhang, Yaxi Liu, Xiaoying Fan, Shanshan Ai, Yingjie Luo, Xin Li, Sai Luo, Hui Zheng, Yanzhu Yue, Zai Chang, Zhongzhou Yang, Fuchou Tang, Aibin He, Xiaohua Shen

## Abstract

Rigorous exploration and dissection of potential actions and effects of long noncoding RNA (lncRNA) in animals remain challenging. Here using multiple knockout mouse models and single- cell RNA sequencing, we demonstrate that the divergent lncRNA *Hand2as* has a key, complex modulatory effect on the expression of its neighboring gene *HAND2* and subsequently on heart development and function, largely independent of *Hand2as* transcription and transcripts. Full-length deletion of *Hand2as* in mouse causes moderate yet prevalent upregulation of *HAND2* in hundreds of cardiac cells, resulting in profound biological consequences, including dysregulated cardiac gene programs, congenital heart defects and perinatal lethality. We propose a *cis*-functional role for the *Hand2as* locus in dampening *HAND2* expression to restrain cardiomyocyte proliferation, thereby orchestrating a balanced development of cardiac cell lineages. This study highlights the need for complementary genetic and single-cell approaches to delineate the function and primary molecular effects of an lncRNA in animals.

**Impact statement:** The long noncoding RNA *Hand2as* critically controls the precise expression of its neighboring gene *HAND2*, thereby balancing cardiac lineages and expression programs that are essential for heart development and function.

## Introduction

Long noncoding RNAs (lncRNAs) have been implicated as an important layer of regulatory information in fine-tuning the spatiotemporal expression of pleiotropic developmental loci in their chromatin neighborhood, thereby modulating cell fate determination in various biological processes (Han et al., 2018; Luo et al., 2016; Morris and Mattick, 2014; Pauli et al., 2011; Ponjavic et al., 2009; Yin et al., 2015). Heart formation is tightly regulated during mouse embryogenesis and involves restriction of mesodermal precursor cells to the cardiac lineage and the subsequent formation of a primitive heart tube, which, in turn, undergoes looping, formation of the outflow tract and atrial and ventricular cavities, and septation to form the mature four-chambered heart (Bruneau, 2008; Olson and Schneider, 2003). Proper commitment of cardiac lineages during this complex process is required for normal development and function of the heart (Brade et al., 2013). Several lncRNAs have been reported to have roles in regulating heart development and function. For example, depletion of *Chast/Wisper* and overexpression of *Mhrt*, *Tincr*, or *Carel* protected the heart from hypertrophy in response to pressure overload following transverse aortic constriction surgery (Cai et al., 2018; Han et al., 2014b; Micheletti et al., 2017; Shao et al., 2017; Viereck et al., 2016). Inhibition of *Fendrr* led to embryonic lethality around E13.5 with cardiac hypoplasia (Grote et al., 2013).

The lncRNA *Hand2as* (also named *Uph* or *lncHand2*) is divergently positioned at −123 bp upstream of the transcription start site (TSS) of *HAND2* (Anderson et al., 2016; Wang et al., 2018). HAND2, a transcription factor that promotes ventricular cardiomyocyte expansion and cardiac reprogramming, is a critical regulator of embryonic heart development (McFadden et al., 2005; Song et al., 2012; Srivastava et al., 1997). RNA *in situ* hybridization analysis of mouse embryos reveals that cardiac expression of *HAND2* is initially detected in the cardiac crescent at E7.75 and continues throughout the linear heart tube at E8.5, and thereafter is specifically enhanced in the developing right ventricle (RV) and outflow tract (OFT), with lower levels of expression in the atrial and left ventricular chamber (Srivastava et al., 1997; Tamura et al., 2014). This pattern persists through E9.5-E10.0, after which *HAND2* expression is downregulated in the cardiac mesoderm but is maintained in the neural crest-derived aortic arch arteries (Srivastava et al., 1997; Tamura et al., 2014).

Precise expression of *HAND2* is essential for normal heart morphogenesis and function, and is tightly regulated at the transcriptional level by a network of cardiac transcription factors and upstream enhancers, and at the post-transcriptional level by microRNAs (Bruneau, 2005; Dirkx et al., 2013; McFadden et al., 2005; McFadden et al., 2000; Zhao et al., 2007; Zhao et al., 2005). Constitutive *HAND2* knockout (KO) in mice caused right ventricle hypoplasia and embryonic lethality at E10.5 (Srivastava et al., 1997). Conditional ablation of *HAND2* in specific sets of cardiac cells led to embryonic lethality at various stages prior to embryonic day E15.5 (summarized in Supplementary file 1) (Holler et al., 2010; Morikawa and Cserjesi, 2008; Morikawa et al., 2007; Tsuchihashi et al., 2011; VanDusen et al., 2014). Overexpression of *HAND2* in transgenic mouse models also led to heart development defects and malfunctions (Supplementary file 1). For example, embryonic overexpression of *HAND2* driven by the β-myosin heavy chain (*MYH7*) promoter prevented the formation of the interventricular septum in embryos (Togi et al., 2006), while overexpression driven by the α-myosin heavy chain (*MYH6)* promoter resulted in pathological myocardial hypertrophy and heart failure in adult hearts (Dirkx et al., 2013).

In a polyA-knockin (KI) mouse model of *Uph*/*Hand2as* reported previously, termination of transcription by insertion of a triple polyadenylation (polyA) stop sequence into intron 1 of *Uph* (- 644 bp upstream of the *HAND2* TSS) abolished *HAND2* expression and led to failed right ventricle formation and lethality at E10.5, partially phenocopying *HAND2* KO mice (Anderson et al., 2016). In addition, compound heterozygous *Uph* and *HAND2* mutant embryos lack *HAND2* expression and recapitulate the *HAND2* KO phenotype (Anderson et al., 2016). It was concluded that transcription of *Uph*/*Hand2as* is an essential switch for the activation of *HAND2* and the onset of heart morphogenesis (Anderson et al., 2016). However, the functional role of *Hand2as* transcripts and the *Hand2as* DNA sequences in the heart remains elusive.

To delineate the role of *Hand2as* in heart development and function, we generated three deletion alleles of *Hand2as* in mouse (Han et al., 2018). Full-length deletion of the entire *Hand2as* sequence (*Hand2as*^F/F^ KO) led to dysregulated cardiac gene expression programs, ventricular septal defects and heart hypoplasia and perinatal death, reminiscent of congenital heart diseases. A short distal deletion at the 3’ end of the *Hand2as* locus (*Hand2as*^D/D^ KO) caused severe contraction defects in adult heart that progressively worsened with increasing age. In comparison, short deletion of the 5’ promoter and exons of *Hand2as* (*Hand2as*^P/P^ KO) effectively diminished *Hand2as* expression, but failed to produce discernable heart phenotypes in either embryos or adults. These results indicate that the *Hand2as* DNA locus, rather than its transcription/transcripts, primarily controls heart development and function. To our surprise, cardiac expression of *HAND2* was sustained in all three *Hand2as* KO mouse models we generated, in sharp contrast to the abolished expression of *HAND2* in the *Uph*/*Hand2as* polyA KI embryos (Anderson et al., 2016). Importantly, single-cell transcriptomic analysis revealed subtle yet prevalent upregulation of *HAND2* and concordant global gene expression changes in subsets of cardiac cells of *Hand2as*^F/F^ embryos lacking the entire *Hand2as* DNA sequence. Altogether, these results illustrate a fine-tuning yet critical role for the lncRNA *Hand2as* locus in restricting the precise, spatial expression of *HAND2*, through which *Hand2as* modulates cardiac lineage development and heart function. This study reveals the unexpected complexity of lncRNA function *in vivo*, and also emphasize the usage of complementary genetic and single-cell approaches to delineate the primary molecular effects and elucidate physiological functions of an lncRNA in animals.

## Results

### *Hand2as* transcripts are dispensable for heart development

*Hand2as* and *HAND2* are divergently transcribed from the shared core promoter sequences, and are highly enriched in the heart compared to other tissue types analyzed (Figure 1A, Figure 1-figure supplement 1A-B). Interesting, within the heart, *Hand2as* and *HAND2* exhibit an inverse expression pattern during embryonic heart development and postnatal growth (Figure 1-figure supplement 1C).

**Figure 1.**
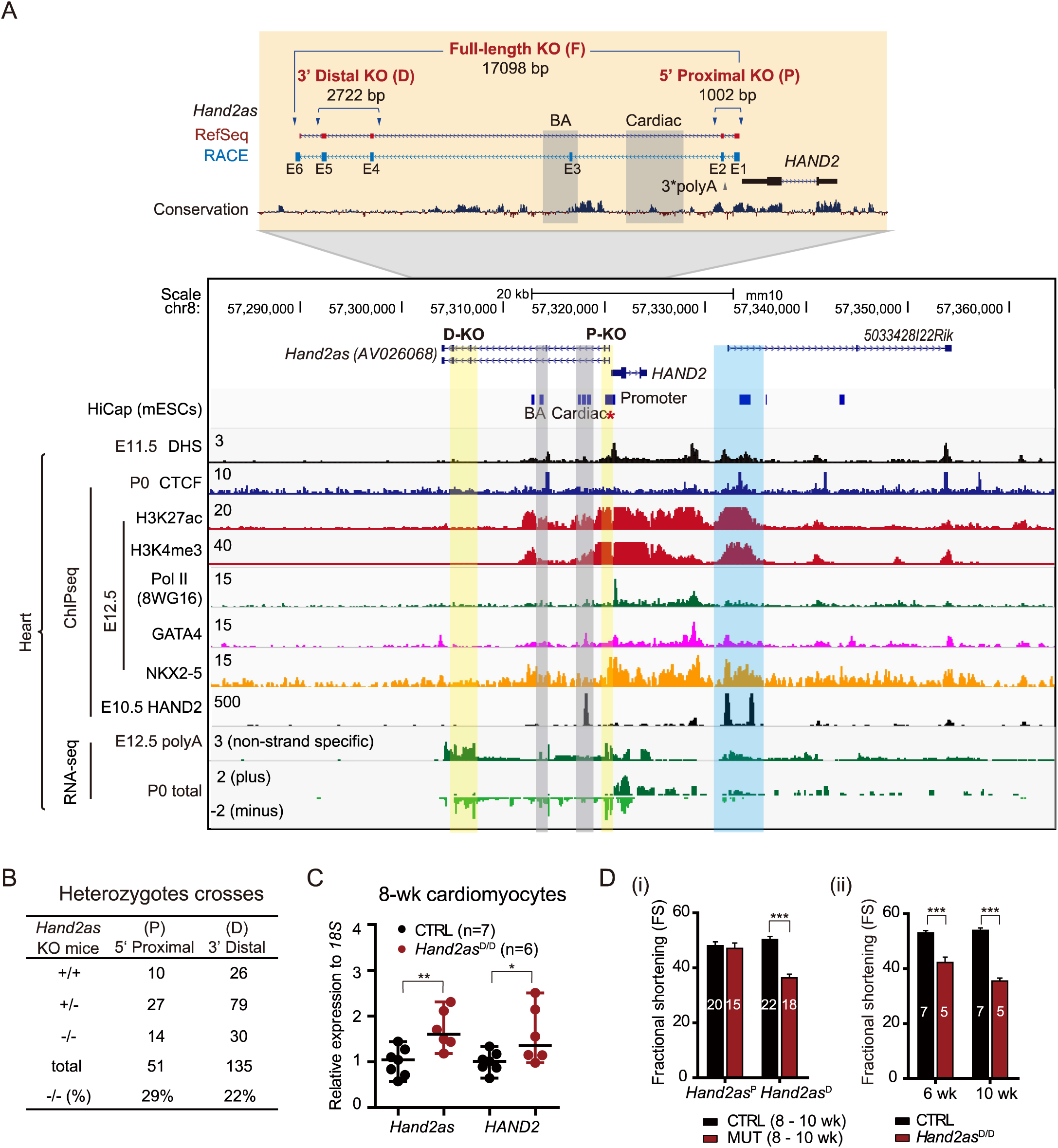
*Hand2as* transcripts are dispensable for cardiac development. *(A)Hand2as* knockout strategies for 5’ proximal KO (P), 3’ distal KO (D) and full-length KO (F). The location of the 3*polyA insertion by Anderson *et al*. is shown, along with the full *Hand2as* transcript characterized by Rapid Amplification of cDNA Ends (RACE). The exons of *Hand2as* are numbered from E1 to E6. The brackets show deleted regions. Arrowheads indicate sgRNA targeting sites. The UCSC genome browser view shows conservation, DHS, potential chromatin interaction (HiCap, CTCF ChIP), histone modification (H3K27ac, H3K4me3), Pol II binding signal, transcription factor (GATA4, NKX2-5, HAND2) binding signals, and RNA signal of the *Hand2as*/*HAND2* locus, in a cell line or tissues at various developmental stages as indicated. Red asterisk, promoter used in HiCap; blue shading, putative enhancers; yellow shading, deleted regions; gray shading, the branchial arch (BA) enhancer and the cardiac enhancer. (B) Survival analysis of *Hand2as* KO lines (*Hand2as*^P^ and *Hand2as*^D^) determined by heterozygote intercrosses. Total, number of analyzed mice; -/- (%), percentage of -/- (homozygotes) out of the total number of analyzed mice. (C) RT-qPCR analysis of *Hand2as* (primers P) and *HAND2* (CDS) in 8-wk cardiomyocytes from CTRL (*Hand2as*^+/+, D/+^) and *Hand2as*^D/D^. The y axis shows expression relative to *18S* (normalized to CTRL). Data are shown as median with range. *, p<0.05; **, p<0.01. (D) Echocardiographic measurement of fractional shortening (FS) to evaluate cardiac function in *Hand2as*^P/P^ and *Hand2as*^D/D^ mice of 8–10 weeks old and wild-type or heterozygous littermates (i), and progressive cardiac function of *Hand2as*^D/D^ mice at 6 weeks and 10 weeks old compared to wild- type or heterozygous littermates (ii). Data are shown as mean ± SEM (n is indicated on each column). ***, p<0.001. n, number of analyzed mice. See also Figure 1-figure supplement 1 and Supplementary files 2, 3 and 9.

The DNA sequence of *Hand2as* is 17 kb in length and encompasses a super-enhancer element, and branchial arch (BA) and cardiac enhancers annotated previously (Figure 1A) (McFadden et al., 2000; Yanagisawa et al., 2003). In E12.5 embryonic hearts, the *Hand2as* locus as well as *HAND2* and its downstream regions harbor multiple DNase I hypersensitive sites (DHS), and show strong binding signals of active histone H3K4me3 and H3K27ac marks (H3 Lys4 tri-methylation and Lys27 acetylation, respectively), and RNA polymerase II and master transcription regulators of cardiac development, including GATA4, NKX2-5 and HAND2 (E10.5) itself (Figure 1A) (He et al., 2014; Laurent et al., 2017; Ye et al., 2015; Yue et al., 2014). This suggests a possible involvement of multiple enhancers in regulating *HAND2* expression. In support of this notion, we found that in mouse embryonic stem cells (ESCs), the *HAND2* promoter interacts with two known upstream enhancers (BA and cardiac) in the *Hand2as* locus, and also with multiple downstream DNA elements embedded in the lncRNA *5033428I22Rik* locus (a strong interaction within the first intron of *5033428I22Rik* is marked by blue shading in Figure 1A), as shown by HiCap, a genome-wide promoter capture method to detect chromatin contacts (Sahlen et al., 2015).

We set out to dissect the function of *Hand2as* in the heart. To remove *Hand2as* transcription/transcripts with minimal manipulation of the genome, we first generated two mouse models carrying short genomic deletions at the 5’ or 3’ end of the *Hand2as* locus (Figure 1A) (Han et al., 2018). These deletions do not affect the two known enhancers for cardiac and branchial arch expression of *HAND2* (McFadden et al., 2000; Yanagisawa et al., 2003). In the 5’ proximal knockout allele (*Hand2as*^P^ KO), we deleted a 1-kb DNA sequence covering the core promoter and the first two exons of *Hand2as* (Figure 1A) (Han et al., 2018). The deletion starts at −61 bp and −62 bp upstream of the TSSs of *Hand2as* and *HAND2* respectively. To avoid any direct effect of promoter alteration on *HAND2* expression, we generated a 3’ distal knockout allele (*Hand2as*^D^ KO) by deleting a 2.7-kb DNA sequence which spans exons 4 and 5 of *Hand2as* and is located 13-kb upstream of the *HAND2* TSS (Figure 1A) (Han et al., 2018).

Both *Hand2as*^P/P^ and *Hand2as*^D/D^ mice from heterozygotes crosses were born at the expected Mendelian ratio and had no overt morphological defects in the heart (Figure 1B, Figure 1-figure supplement 1A; data not shown). Compared to heterozygous littermates, *Hand2as*^P/P^ mutant mice showed residual expression (10~17%) of a truncated *Hand2as* RNA which lacks the first two exons and represents 64% of the full-length transcript, in hearts from E12.5 embryos or 8-week old adults (Figure 1-figure supplement 1A and 1D). *Hand2as*^P/P^ KO mice therefore provide a partial loss-of- function model. However, cardiac expression of *HAND2* was not altered in *Hand2as*^P/P^ mice (Figure 1-figure supplement 1A and 1D).

In comparison, *Hand2as*^D/D^ mice expressed a mutant *Hand2as* RNA that lacks exons 4 and 5 with 67% of its sequence remaining (Figure 1-figure supplement 1A). Embryonic expression of *HAND2* in *Hand2as*^D/D^ hearts was not affected (data not shown). However, *Hand2as*^D/D^ adult cardiomyocytes showed moderate but significant increases of both *Hand2as* and *HAND2* transcripts (~53% and ~34% up, respectively) (Figure 1C). It was reported that aberrant upregulation of *HAND2* in the postnatal heart contributes to pathological hypertrophy (Dirkx et al., 2013). Interestingly, *Hand2as*^D/D^, but not *Hand2as*^P/P^ mice, progressively developed heart contraction defects at 6-10 weeks old, with a 10~30% decrease in fractional shortening (Figure 1D). Consistent with the cardiac contraction defect, many genes involved in heart development, cardiac muscle contraction, and the cell cycle were dysregulated in *Hand2as*^D/D^ cardiomyocytes (Figure 1-figure supplement 1E-F; Supplementary files 2 and 3). These results suggested that the *Hand2as* locus may exert a complex, pleiotropic influence on *HAND2* expression and heart physiology. However, the lack of heart phenotype in *Hand2as*^P/P^ mice indicates that *Hand2as* transcripts and perhaps its transcription may be largely dispensable for cardiac expression of *HAND2* and embryonic heart development.

### Deletion of the entire *Hand2as* locus causes congenital heart defects and perinatal lethality

Next, to rule out the possibility that residual activities of *Hand2as* might promote *HAND2* expression and heart morphogenesis in the promoter and distal KO mouse models, we deleted a 17- kb sequence covering the entire *Hand2as* genomic region to completely eliminate *Hand2as* expression (Figure 1A) (Han et al., 2018). The deletion starts at −59 bp and −64 bp upstream of *Hand2as* and *HAND2* TSSs, respectively, and encompasses the super-enhancer and two known enhancers of *HAND2* expression (McFadden et al., 2000; Yanagisawa et al., 2003) (Figure 1A). This mutant allele is designated as *Hand2as* full-length knockout (*Hand2as*^F^ KO).

Heterozygous *Hand2as*^F/+^ intercrosses failed to produce viable homozygous offspring (0 out of 77 pups) at the weaning stage (Figure 2A, Figure 1-figure supplement 1A). Viable *Hand2as*^F/F^ embryos were observed during mid-gestation at the expected Mendelian frequency until E16.5, but thereafter, death occurred at varying times, ranging from E16.5 to just after birth (Figure 2A). *Hand2as*^F/F^ newborns (7 out of 41 at P0) became cyanotic and invariably died shortly after birth (Figure 2B). Gross morphological examination of hearts revealed abnormal blood coagulation and fatal thrombosis in *Hand2as*^F/F^ newborns (Figure 2C), indicating heart failure in response to increased demand for cardiac output upon birth. To uncover the heart defect causing the perinatal lethality, *Hand2as*^F/F^ late-gestation embryos (E16.5) were subjected to necropsy and histological examination. Macroscopically, the most severe phenotype in homozygous embryonic hearts is the presence of ventricular septal defects (8 out of 12) (Figure 2D). These lesions may cause blood to leak from the left ventricle into the right ventricle and then a right-to-left shunt, consequently leading to immediate cyanosis and death (Minette and Sahn, 2006). Right ventricular (RV) hypoplasia was also frequently observed with significantly decreased chamber volume (10 out of 12) and slightly reduced thickness of the compact myocardium of the right ventricle (3 out of 12) in *Hand2as*^F/F^ mutant hearts (Figure 2D). These defects are reminiscent of congenital heart diseases, and provide a morphological explanation for the heart failure of *Hand2as*^F/F^ mice in response to cardiac stress at birth.

**Figure 2.**
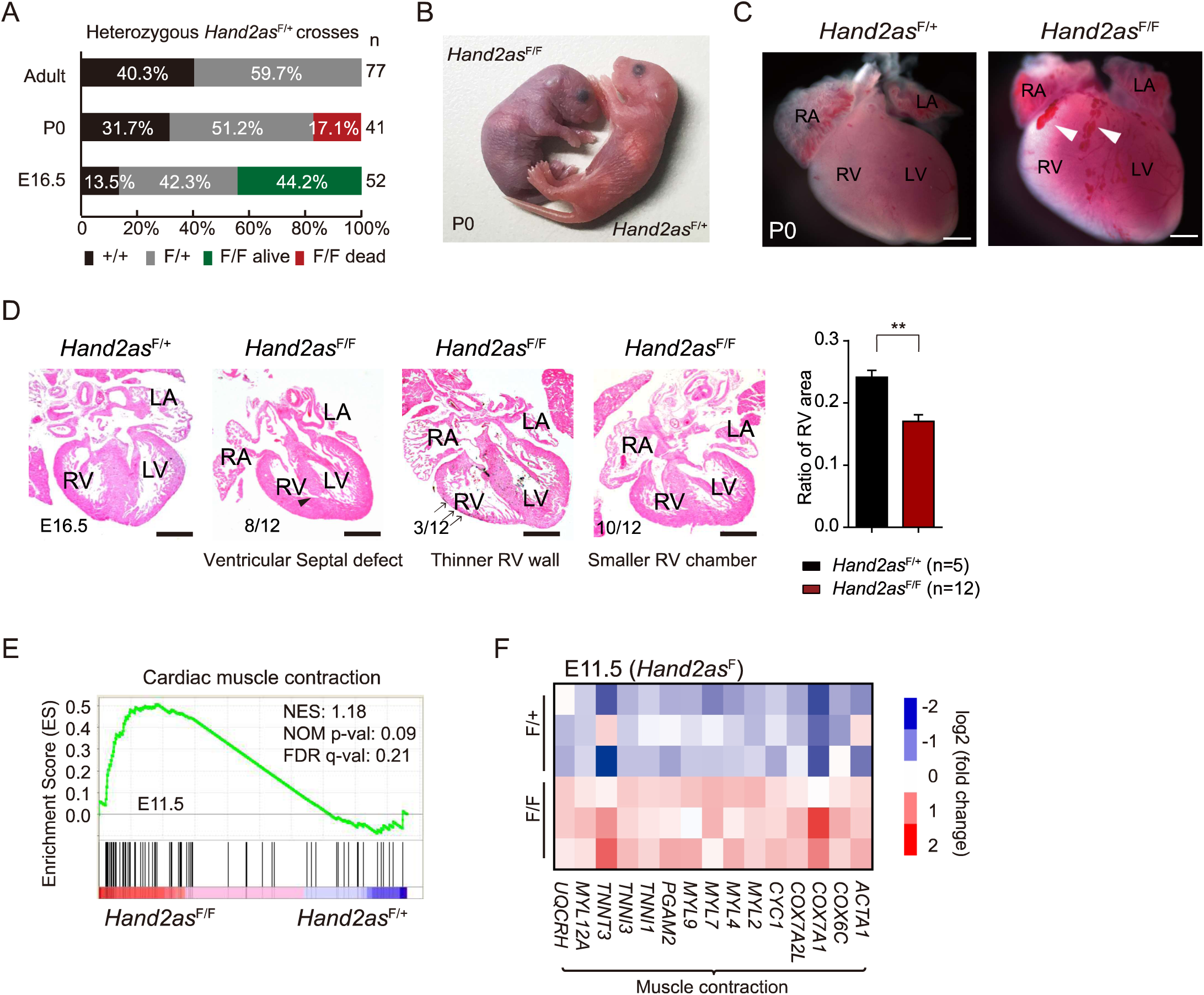
Deletion of the entire *Hand2as* locus causes congenital heart defects and perinatal lethality. (A) Survival analysis of *Hand2as*^F/+^ intercrosses at three stages (E16.5, P0 and adult). (B) *Hand2as*^F/+^ and *Hand2as*^F/F^ newborns. *Hand2as*^F/F^ mice turned cyanotic, demonstrating poor circulation compared to the ruddy *Hand2as*^F/+^ mice (*Hand2as*^F/F^, n=3; *Hand2as*^F/+^, n=7). (C) Gross morphological examination of P0 hearts of *Hand2as*^F/+^ and *Hand2as*^F/F^ mice reveals abnormal blood coagulation and fatal thrombosis (arrowheads) in *Hand2as*^F/F^ newborns. (D) Hematoxylin-and-eosin staining of transverse sections of E16.5 hearts (left) and morphometric analysis of the ratio of right ventricle area (normalized to whole ventricle area). Number of hearts with a defect similar to the representative picture is shown in the bottom left corner. The arrowhead indicates the lesion in the ventricular septum. Arrows indicate the right ventricular compact myocardium. Scale bar, 250 µm. Data are shown as mean ± SEM. **, p<0.01. (E) Upregulation of genes related to cardiac muscle contraction (KEGG PATHWAY: mmu04260) in *Hand2as*^F/F^ E11.5 hearts by Gene Set Enrichment Analysis (GSEA) (n=3 for each genotype). (F) Heatmap showing representative genes that are dysregulated in E11.5 hearts of *Hand2as*^F/F^compared to *Hand2as*^F/+^ (n=3 for each genotype). n, number of analyzed mice; RV, right ventricle; LV, left ventricle; RA, right atrium; LA, left atrium; NES, Normalized Enrichment Score. See also Figure 2-figure supplement 1 and Supplementary files 2, 4, 5 and 9.

Notably, *Hand2as*^F/F^ late-gestation embryos and newborns had cleft palate (Figure 2-figure supplement 1A), resembling the craniofacial defects observed in branchial arch enhancer KO mice, which reportedly, failed to suckle and died with an empty stomach 24 hours after birth (Yanagisawa et al., 2003). As *Hand2as*^F/F^ pups died much earlier, within hours after birth, we reasoned that the suckling defect is not the cause of death in these animals. In addition, we found no gross abnormalities in other organs, including aortic arch arteries, liver and lung (Figure 2-figure supplement 1B-C). Moreover, *Hand2as*^F/F^ pups showed normal floating lungs in a buoyancy test (data not shown), excluding the possibility of respiratory failure as the cause for their immediate death upon birth.

To reveal transcriptional changes that underlie the morphological defects and perinatal lethality, we performed RNA-seq analysis of embryonic hearts or ventricles isolated from littermates from *Hand2as*^F/+^ intercrosses. Interestingly, a subset of gene programs pertaining to cardiac muscle contraction, such as *ACTA1*, *COX6C* and *MYL2* were upregulated in *Hand2as*^F/F^ embryonic hearts at E11.5, implying abnormally increased cardiac myogenesis in the mutant heart (Figure 2E-F; Supplementary files 2 and 4). Further transcriptome analysis of E16.5 ventricles also revealed significant upregulation of genes related to hypertrophic cardiomyopathy in *Hand2as*^F/F^ embryos (Figure 2-figure supplement 1D; Supplementary files 2 and 5). The transcriptomic defects offer a molecular explanation for heart morphological defects and function failure, which are most likely the cause of the perinatal death of *Hand2as*^F/F^ newborns.

### Sustained *HAND2* expression in *Hand2as*^F/F^ embryos

To study the direct effect of *Hand2as* deletion on *HAND2* expression, we first confirmed the complete absence of *Hand2as* transcripts in *Hand2as*^F/F^ mutant embryos by RNA-seq and RT-qPCR (reverse transcription and quantitative PCR) (Figure 3-figure supplement 1A; data not shown). In addition, we did not observe any RNA signals or transcripts downstream of the *Hand2as* locus (data not shown). Thus, *Hand2as*^F/F^ KO mice provided a complete loss-of-function model, in which the transcription, transcripts and DNA sequences of *Hand2as* were simultaneously removed. However, to our surprise, the levels of cardiac *HAND2* transcripts were not lost in *Hand2as*^F/F^ mutant embryos. The coding sequence (CDS) of *HAND2* showed comparable expression between homozygous and heterozygous littermates throughout heart morphogenesis from E9.5 to E16.5 (Figure 3A-B).

**Figure 3.**
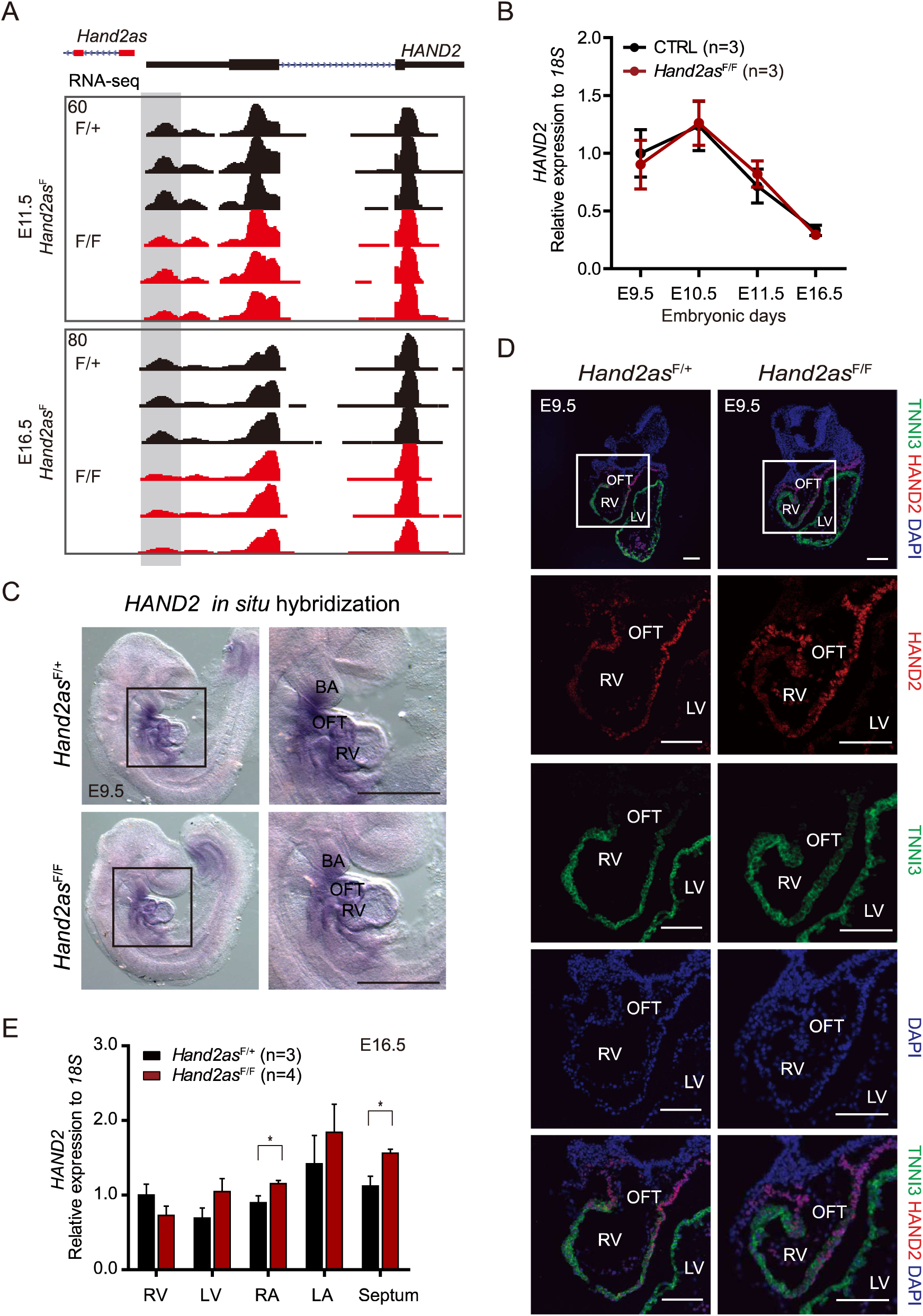
Sustained *HAND2* expression in *Hand2as*^F/F^ embryos. (A) The genome browser view shows expression (RNA-seq signal) of *HAND2* in E11.5 hearts (top) and E16.5 ventricles (bottom) from *Hand2as*^F/F^ mutants compared to *Hand2as*^F/+^ littermates (n=3 for each genotype). The gray shading indicates the *HAND*2 5’ UTR region in which the signal is downregulated. (B) RT-qPCR analysis of *HAND2* (coding sequence) in embryonic hearts at various stages from CTRL (*Hand2as*^+/+, F/+^) and *Hand2as*^F/F^. The y axis shows expression relative to *18S* (normalized to *HAND2* expression in CTRL embryos at E9.5). Data are shown as mean ± SEM. (C) Whole mount *in situ* hybridization of *Hand2as*^F/F^ and *Hand2as*^F/+^ embryos at E9.5 shows *HAND2* RNA expression in heart and branchial arch. Zoomed-in images of the boxed areas are shown on the right (*Hand2as*^F/F^, n=2; *Hand2as*^F/+^, n=3). Scale bar, 500 μm. (D) Immunofluorescence staining of HAND2 and TNNI3 (marker of cardiomyocytes) in *Hand2as*^F/+^ and *Hand2as*^F/F^ embryonic hearts at E9.5. Scale bar, 100 μm. The four lower panels show zoomed- in images of the boxed areas in the top images (n=3 for each genotype). (E) RT-qPCR analysis of *HAND2* (coding sequence) in dissected E16.5 embryonic hearts from *Hand2as*^F/+^ and *Hand2as*^F/F^. The y axis shows expression relative to *18S* (normalized to *HAND2* expression in RV of *Hand2as*^F/+^ embryos). Data are shown as mean ± SEM. *, p<0.05. n, number of analyzed mice; RV, right ventricle; LV, left ventricle; RA, right atrium; LA, left atrium; OFT, outflow tract; BA, branchial arch. See also Figure 3-figure supplement 1 and Supplementary file 9.

To confirm this finding, we performed RNA *in situ* hybridization (ISH) analysis of E9.5 embryos and found similar distribution and expression patterns of *HAND2* mRNA between homozygous and heterozygous littermates (Figure 3C). Next we analyzed HAND2 protein levels by immunostaining analysis of transverse sections of E9.5 embryonic hearts. Again, comparable levels of HAND2 protein were detected in mutant embryos (Figure 3D, Figure 3-figure supplement 1B). We noted a 1-2-fold reduction in *HAND2* RNA signals that fall into its 5’ untranslated region (UTR) in *Hand2as*^F/F^ embryonic hearts, despite unchanged expression in the CDS of *HAND2* (Figure 3A, Figure 3-figure supplement 1C). Shortening of the *HAND2* 5’ UTR might promote translation of HAND2 protein, as demonstrated in cultured cells (Figure 3-figure supplement 1D) (Curtis et al., 1995; Leppek et al., 2018). Nevertheless, the overall expression of *HAND2* at both the RNA and protein levels was sustained in complete absence of *Hand2as*.

The mature four-chamber heart of an E16.5 embryo can be experimentally dissected into distinct compartments, in which *HAND2* transcripts can then be analyzed by RT-qPCR. Interestingly, except for an insignificant and slight decrease in mutant right ventricles, *HAND2* expression showed a tendency to be upregulated in all other compartments of E16.5 *Hand2as*^F/F^ hearts, compared to those of heterozygous littermates (Figure 3E, Figure 3-figure supplement 1E). In particular, *HAND2* expression in the mutants was significantly increased by ~40% in the septum and ~24% in the right atrium (RA) (Figure 3E, Figure 3-figure supplement 1E). This subtle yet significant upregulation of *HAND2* in specific regions of mutant hearts suggests that the *Hand2as* locus might be involved in controlling the spatial expression of *HAND2* during heart formation.

### Single-cell transcriptomic profiling reveals four cardiac cell types

To reveal subtle alterations that are not readily detectable in population-based analysis due to the averaged expression of mixed cells, we performed high-throughput single-cell RNA-seq analysis of E11.5 embryonic hearts isolated from *Hand2as*^F/F^ and wild-type littermates (Figure 4A). To delineate the primary transcriptional effects of *Hand2as* deletion, we chose to analyze embryos at E11.5, which is also the most convenient early time point that we could experimentally isolate enough cells for 10X Genomics cell sorting and library construction. After removing sequencing reads from hematopoietic cells, we obtained expression profiles for a total of 3,600 cardiac cells, including 2,108 for *Hand2as*^F/F^ and 1,492 for the wild type, with an average of 0.06 million reads/~13,000 Unique Molecule Identifier (UMI) counts per cell and a median level of 3,492 expressed genes per cell (transcripts per million [TPM] > 0) (Figure 4A-B; Supplementary file 6).

**Figure 4.**
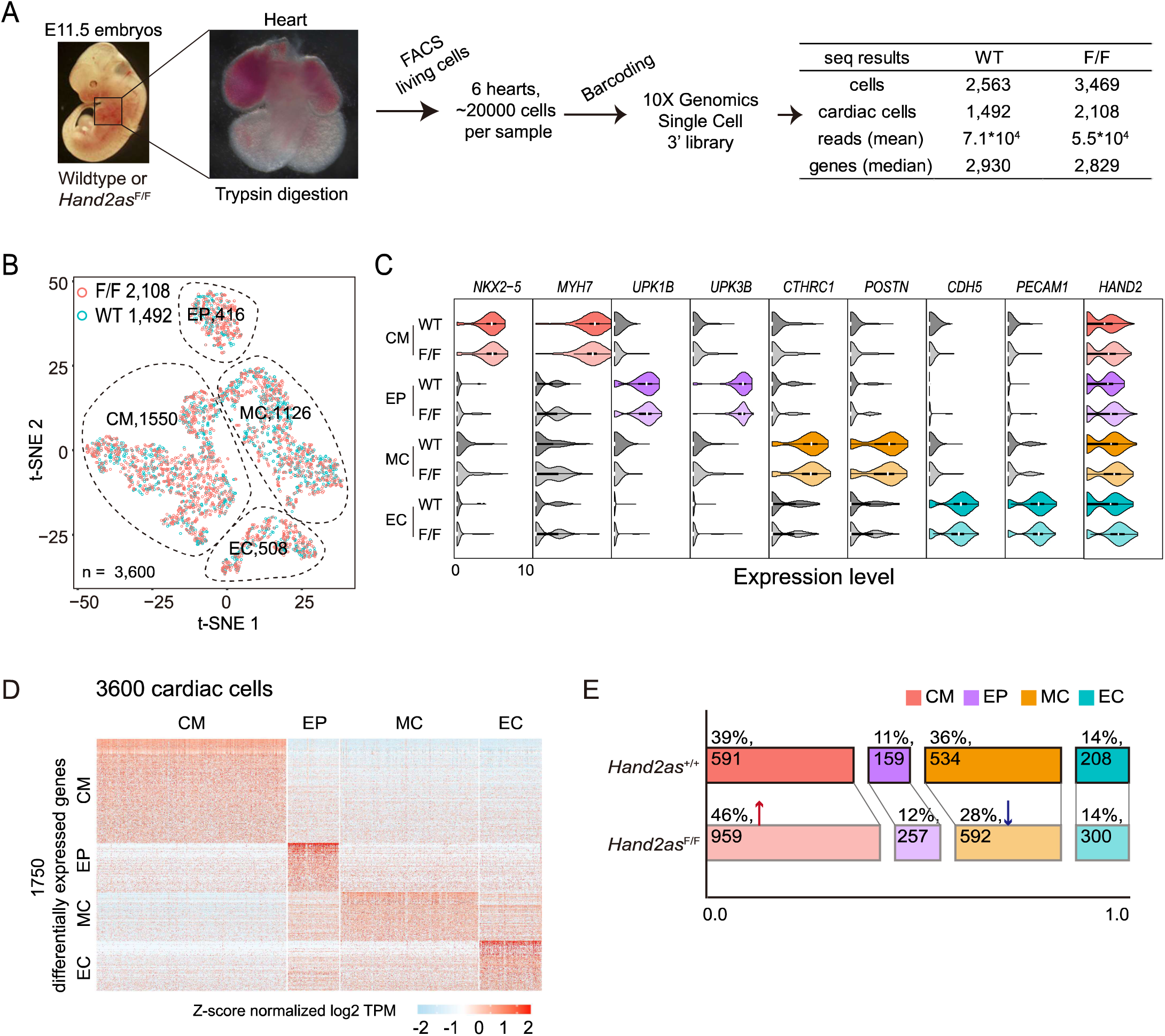
Single-cell transcriptomic profiling reveals four cardiac cell types. (A) Flow diagram and sequencing results of transcriptomic profiling at single-cell resolution using embryonic hearts from E11.5 *Hand2as*^F/F^ and wild-type embryos. (B) Two-dimensional t-SNE visualization of graph-based clustering carried out on *Hand2as*^F/F^ and wild-type embryonic hearts. The dataset comprises a total of 3600 cardiac single-cell transcriptomes. Hematopoietic cells and platelets were removed. The 3600 single-cell transcriptomes were divided into 4 subsets, encircled by dashed lines: CM, cardiomyocyte; EP, epicardial cell; MC, mesenchymal cell; EC, endothelial cell. The number of cells of each type is indicated. (C) Violin plots of gene expression (log2 (TPM/10+1)) for representative markers of each cell type. (D) Heatmap shows Z-score normalized expression of differentially expressed genes (1750) of the four cell types shown in panel B. Rows and columns represent genes and single cells, respectively. (E) Bar charts show distribution of cardiac cells in *Hand2as*^F/F^ and wild-type embryonic hearts. Red arrow, increase of CMs; blue arrow, decrease of MCs. Percentages and cell numbers of each cell type in *Hand2as*^F/F^ or wild-type embryonic hearts are indicated. TPM, Transcripts Per Million; t-SNE, t-distributed stochastic neighbor embedding. See also Supplementary files 6 and 7.

Classification using the t-distributed stochastic neighbor embedding method (t-SNE) revealed four well-separated clusters of cells that differentially express distinct marker genes (Figure 4B-D; Supplementary files 6 and 7). About 43% (1550) of cells are cardiomyocytes (CMs), which specifically express *NKX2-5* and *MYH7* (Figure 4B-C; Supplementary files 6 and 7). About 31% (1126) are mesenchymal cells (MCs) expressing *CTHRC1* and *POSTN*. About 12% (416) of cells are epicardial cells (EPs) marked by membrane proteins *UPK1B* and *UPK3B* and ~14% (508) are endothelial cells (ECs) expressing *CDH5* and *PECAM1* (Figure 4B-C; Supplementary files 6 and 7) (Li et al., 2016). Interestingly, altered proportions of cardiomyocytes and mesenchymal cells were observed in *Hand2as*^F/F^ embryonic hearts compared to the wild-type littermates (Figure 4E). Mutant CMs and MCs increased ~7% and decreased ~8%, respectively, while ratios of mutant EPs and ECs remained similar (Figure 4E). Hundreds of single-cell profiles obtained for each cell type thus provided large numbers of cells for in-depth statistical comparisons of gene expression changes between mutant and wild-type hearts.

### Upregulation of *HAND2* and nearby genes in *Hand2as*^F/F^ embryos

*HAND2* is ubiquitously expressed (TPM>0) in 60% of cardiac cells at E11.5 (Figure 5A). This observation was consistent with previous reports of robust expression of HAND2 detected in many types of cardiac cells using RNA ISH and immunostaining (Laurent et al., 2017; VanDusen and Firulli, 2012). Interestingly, the *Hand2as*^F/F^ hearts appeared to have many more cells (70%) expressing *HAND2*, representing a 10% increase of *HAND2*-positive populations (Figure 5A). The observed increase of mutant CMs mainly resulted from an increase of *HAND2*-positive, but not negative, cardiomyocytes (*p* = 2E-04) (Figure 5B). In comparison, the observed decrease of mutant MCs mainly resulted from a decrease of *HAND2*-negative mesenchymal cells (*p* = 6E-04) (Figure 5B). Thus, the opposing changes in the percentages of mutant CMs and MCs appeared to be positively correlated with changes of *HAND2* expression in these populations.

Although the overall percentage of ECs remained the same, mutant ECs exhibited a significant increase in *HAND2*-positive cells but a decrease in *HAND2*-negative cells (*p* = 4E-09) (Figure 5B). In mutant EPs, *HAND2*-positive cells also increased ~8% (Figure 5A). Thus, cell populations expressing *HAND2* increased ~8-24% in all types of cardiac cell in *Hand2as*^F/F^ mutant hearts, with the most significant gain in the ECs (1.4-fold increase from 59% to 83%, *p* = 4E-09) (Figure 5A). Moreover, the median levels of *HAND2* expression went up significantly by 8-12% across four cardiac cell types (Figure 5A). In comparison, other master regulators of cardiac development, such as *GATA6, GATA4* and *NKX2-5*, either showed unaltered or decreased expression in particular cardiac cell types (Figure 5-figure supplement 1A). These results indicate that the subtle yet global upregulation of *HAND2* is specific to the loss of *Hand2as*, rather than sequencing variations between the wild-type and mutant samples.

**Figure 5.**
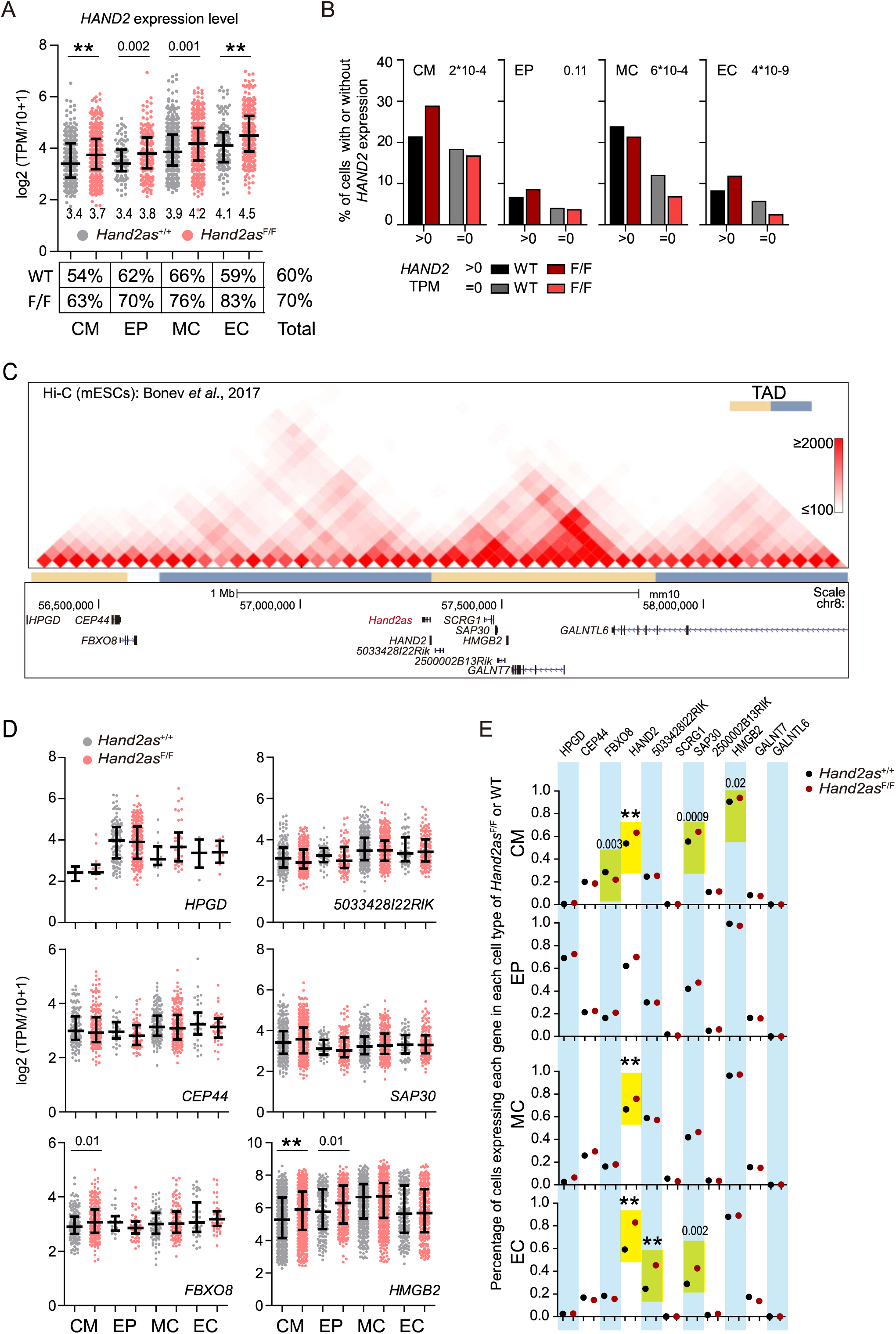
Upregulation of HAND2 and nearby genes in Hand2as^F/F^ embryos. (A) Scatter plots show expression level and frequency of *HAND2* in cardiac cells from *Hand2as*^F/F^ and wild-type embryonic hearts. Data are shown as median and interquartile range. The median value is shown at the bottom of each scatter plot. The *p*-values are indicated at the top: 0.0001<*p*<0.05; **, *p*<0.0001. The percentage of *HAND2*-expressing cells (TPM>0) for each cell type is shown under the plot. (B) Correlation of *HAND2* expression and cardiac cell distribution in *Hand2as*^F/F^ and wild-type embryonic hearts. The y axis shows the percentage of cells with or without *HAND2* expression in *Hand2as*^F/F^ or wild-type embryonic hearts. The *p*-value for Fisher’s test is shown. (C) Genome browser of Hi-C Data shows TADs within ±1-Mb of the *HAND2* TSS, with annotated genes labeled. (D) Scatter plots show expression level of expressed genes (in panel C) in cardiac cells of *Hand2as*^F/F^ and wild-type embryonic hearts. Data are shown as median and interquartile range. The *p*-values are indicated: 0.0001<*p*<0.05; **, *p*<0.0001. A gene is defined as expressed if the corresponding transcripts are detected in more than 20% of cells in at least one cell type. Percentages of cells expressing genes within ±1-Mb of the *HAND2* TSS in each type of cardiac cells. Yellow boxes indicate significant changes of expression frequency between *Hand2as*^F/F^ and wild-type cells. The *p*-values are indicated: 0.0001<*p*<0.05; **, *p*<0.0001.See also Figure 5-figure supplement 1. See also Figure 6-figure supplement 1 and Supplementary files 7 and 8.

Next we sought to determine whether complete removal of *Hand2as* might also affect the expression of other nearby genes. High-order chromatin structure analysis by Hi-C in mouse ESCs (Bonev et al., 2017) showed that the *HAND2* and *Hand2as* locus resides at the boundary of two topologically associating domains (TADs) (Figure 5C). This boundary demarcates an upstream gene desert of ~0.65 Mb in length from a downstream gene-rich region. Among 10 genes located within ±1-Mb genomic regions surrounding the TSSs of *Hand2as* and *HAND2*, we found that 6 genes were expressed in at least one type of cardiac cell, and four of them exhibited very subtle but statistically significant upregulation in the mutant heart (Figure 5D-E).

Of the four genes with significantly altered expression, one (*FBXO8*) lies ~0.75 Mb upstream of *Hand2as* and three (*SAP30, 5033428I22Rik* and *HMGB2*) lie within the same TAD immediately downstream of *HAND2* (Figure 5C). For *SAP30* and *5033428I22Rik* the numbers of cells expressing these genes were significantly upregulated in subsets of mutant cardiac cells (~9-12% increase of *SAP30-*positive CMs and ECs, *p* < 0.002; and ~21% increase of *5033428I22Rik*-positive ECs, *p* = 2E-06) (Figure 5E). *HMGB2* which is highly expressed in all cardiac cells, was specifically upregulated in its transcript abundance in mutant CMs and EPs (*p* < 0.01) (Figure 5D). The median expression level of the upstream gene, *FBXO8* was slightly upregulated (*p* = 0.01) in mutant CMs (Figure 5D). For comparison, we also randomly selected two ubiquitously expressed genes, *EZH2* and *PSMD1*, and found that they showed unaltered expression in both transcript abundance and expression frequency in the mutant hearts (Figure 5-figure supplement 1B). The combined results demonstrate a *cis*-regulatory role for the *Hand2as* locus in dampening the expression of *HAND2* and several neighboring genes in cardiac cells.

### Aberrant cardiac gene programs in *Hand2as*^F/F^ embryos

*HAND2* facilitates cardiomyocyte proliferation and the reprogramming of fibroblasts into functional cardiac-like myocytes *in vitro* and *in vivo* (Song et al., 2012). We next asked how excess amounts of *HAND2* might affect cardiac gene expression programs. Gene ontology (GO) analysis of dysregulated genes showed a significant enrichment of the functional term related to multicellular organism development in all four types of cardiac cell. Interestingly, only non-CMs are specifically enriched in functional terms related to cardiac muscle contraction and heart morphogenesis (Figure 6A; Supplementary file 8). GSEA further revealed global upregulation of muscle contraction genes and downregulation of genes involved in cardiac septum development, both of which were specifically observed in non-CMs of the *Hand2as*^F/F^ mutant, but not in mutant CMs (Figure 6B-C, Figure 6-figure supplement 1; Supplementary file 2). These global, opposing changes in genes related to muscle contraction and cardiac septum development were also confirmed by gene expression of averaged single-cell and bulk RNA-seq of the E11.5 hearts (Figure 6D). Consistent with GSEA, mutant non-CMs exhibited more dramatic expression changes than mutant CMs (Figure 6D).

**Figure 6.**
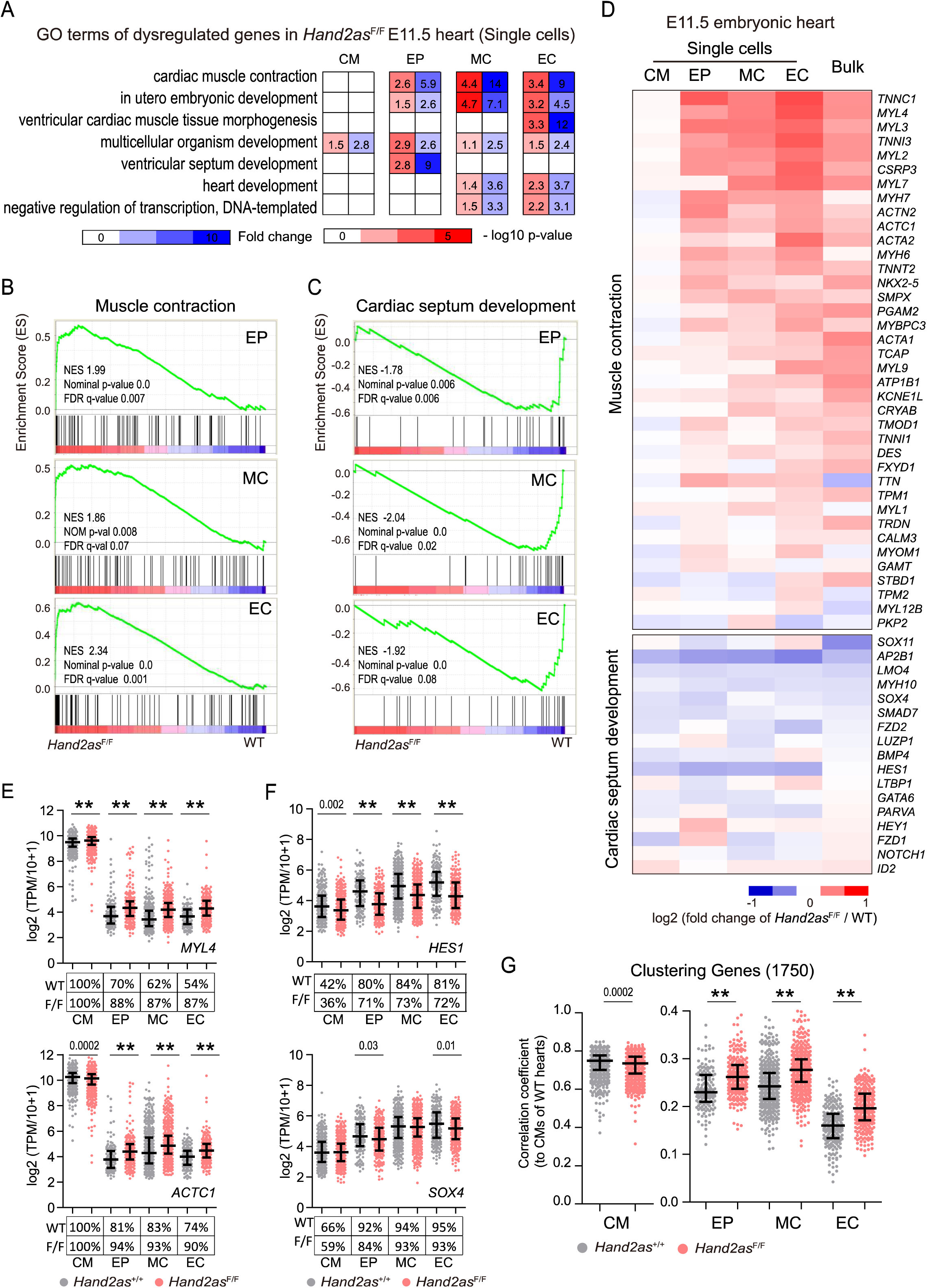
Aberrant cardiac gene programs in *Hand2as*^F/F^ embryos. (A) Heatmap shows enriched Gene Ontology terms (*p*-value <0.01, >5 genes) of dysregulated genes in each cell type. (B-C), GSEA analysis of genes related to muscle contraction (GO: 0006936) (B) and cardiac septum development (GO: 0003279) (C) in non-CMs (D) Heatmap showing fold change of representative genes involved in muscle contraction and cardiac septum development in E11.5 hearts of *Hand2as*^F/F^ compared to wild type. For single-cell RNA-seq, fold change of averaged gene expression in each cell is shown. For bulk RNA-seq of E11.5 embryonic heart, fold change of averaged gene expression from replicas (n=3 for each genotype) is shown. (E-F) Scatter plots show expression level and frequency of *MYL4* and *ACTC1* (E), and *HES1* and *SOX4* (F) in cardiac cells of *Hand2as*^F/F^ and wild-type embryonic hearts. Data are shown as median and interquartile range. The *p*-values are indicated: 0.0001<*p*<0.05; **, *p*<0.0001. Percentages of expressing cells (TPM>0) in each cell type are shown under the plots. (G) Scatter plots show Pearson correlation coefficient for each cell when compared single-cell gene expression profile to the averaged gene expression profile (of 1750 differentially expressed genes shown in Figure 4D) of wild-type CMs. Data are shown as median and interquartile range. The *p*- values are indicated: 0.0001<*p*<0.05; **, *p*<0.0001. See also Figure 6-figure supplement 1 and Supplementary files 7 and 8.

Many CM marker genes regulating muscle contraction and heart development, such as *MYL4*, *ACTC1, TNNC1*, and *TNNT2*, were significantly upregulated in their expression levels and frequencies (Figure 6D-E). In contrast, marker genes enriched in non-CMs, such as *HES1, SOX4* and *FZD2*, which are involved in septum development, were specifically downregulated (Figure 6D and 6F). For example, ratios of *MYL4*-positive cells in mutant hearts were increased 18-33% in non-CMs (*p* < 9E-06), and this was also accompanied by 17-22% increases (*p* < 0.0001) of *MYL4* transcript abundance (Figure 6E). In comparison, 9~11% of non-CMs lost *HES1* expression (*p* < 0.05), and *HES1* RNA abundance went down 12-18% (*p* < 0.0001) in mutant hearts (Figure 6F). These molecular changes explain the ventricular septal defects and ventricle hypoplasia observed in *Hand2as*^F/F^ embryonic hearts. The concurrent upregulation of CM marker genes and downregulation of non-CM marker genes in the non-CM cells of *Hand2as*^F/F^ hearts indicated that mutant non-CMs are aberrantly reprogrammed towards cardiomyocytes.

To further support this finding, we performed correlation analysis to compare the expression similarity of a panel of 1750 differentially expressed genes (DEGs) (Figure 4D) in each of the four cardiac cell types (from wild type and mutant) to those of wild-type CMs. Mutant and wild-type CMs had the highest median levels of correlation coefficient compared to non-CM cells, indicating the robustness of this assay (Figure 6G). Interestingly, compared to their wild-type counterparts, mutant non-CMs (EPs, MCs and ECs) showed significantly higher correlations with cardiomyocytes (Figure 6G). Thus, the gene programs in the non-CMs of *Hand2as*^F/F^ embryos may resemble the gene program in cardiomyocytes (Figure 6G). Our data support a model in which an apparently subtle but global increase in the *HAND2* dose in all cardiac cell types may lead to pronounced changes in cardiac gene programs, and eventually result in morphological and functional abnormalities in *Hand2as*^F/F^ mutant hearts.

## Discussion

Despite extensive reports of lncRNA functions in cell lines, rigorous exploration and dissection of their potential actions and effects in animals are still lacking. Precise expression of *HAND2* is critical for heart formation. Transcription of the divergent lncRNA *Hand2as* was reportedly essential for *HAND2* activation and heart morphogenesis. Using three knockout mouse models, we demonstrate that the *Hand2as* DNA locus, rather than its transcription/transcripts, controls *HAND2* expression and normal heart development and function. Full-length deletion of *Hand2as* led to congenital heart defects and perinatal lethality. Importantly, in embryos lacking the entire *Hand2as* DNA sequence, single-cell transcriptomic analysis of the heart revealed subtle yet prevalent upregulation of *HAND2* and dysregulated cardiac gene expression programs. These results illustrate a critical, fine-tuning function of the lncRNA *Hand2as* locus in accurately controlling the spatial expression of *HAND2*, through which *Hand2as* modulates cardiac lineage development and heart function.

*Hand2as*^P/P^ mice lacked any discernable phenotype in the heart and in animal survival. Consistent with our finding, a recent study of a *Hand2as* (*lncHand2)* mutant mouse model reported that deletion of the first two exons of *Hand2as* (*lncHand2)* did not cause apparent heart abnormality and failed to affect hepatic expression of *HAND2* and other neighboring genes despite the absence of *lncHand2* transcripts in mouse livers (Wang et al., 2018). Although we cannot rule out a potential role for residual (10%) *Hand2as* transcription/RNA in *Hand2as*^P/P^ embryos, it is most likely in this case that disruption of the *Hand2as* DNA locus rather than *Hand2as* transcription/transcripts primarily contributes to heart morphological defects and lethality. Upregulation of *HAND2* and its nearby genes in *Hand2as*^F/F^ mutant hearts mainly resulted from the loss of *cis*-regulatory DNA sequences embedded in the *Hand2as* locus, rather than being a *trans*-acting consequence of developmental defects observed in these mice. The combined results from three *Hand2as* KO mice argue against a switch-like role for *Hand2as* transcription, transcripts and DNA sequences in regulating *HAND2* activation in the heart (Anderson et al., 2016). Instead, the *Hand2as* locus acts as a nudger and tweaker to fine-tune the spatiotemporal expression of cardiac *HAND2* and lineage expression programs. *Hand2as* transcripts might serve as proxy signals for the activity of important regulatory DNA elements for *HAND2* expression (Mowel et al., 2018). However, it remains possible that in other pathological or stress conditions yet to be revealed, *Hand2as* transcription and transcripts may play a role in defining the chromatin environment required for the precise regulation of *HAND2* transcription.

Complete removal of the entire 17-kb *Hand2as* sequence abolishes *Hand2as* transcription and transcripts, but fails to attenuate *HAND2* expression, leading to much weaker cardiac defects and delayed onset of death, in sharp contrast to the abolished expression of *HAND2* and failed heart morphogenesis at E10.5 in *Hand2as/Uph* polyA KI embryos (Anderson et al., 2016). These discrepancies were unexpected, as polyA KI should minimally disrupt the genomic DNA compared to a large deletion. We postulate that the severe phenotypes observed in the polyA KI mice might result from polyA-induced aberrant silencing of the nearby *HAND2* locus that occurs independently of the lncRNA’s function. Because of the close juxtaposition of *Hand2as* and *HAND2*, the proximal insertion of a transcription stop signal immediate upstream (−644 bp) of the *HAND2* TSS might artificially recruit the transcription termination machinery to inhibit the removal of the inhibitory H3K27me3 mark in the *HAND2* promoter of mesodermal precursor cells (Almada et al., 2013), consequently inhibiting *HAND2* activation during the onset of cardiogenesis. One example to support this possibility is the study of the lncRNA *ThymoD* in activating its upstream gene *BCL11b* in developing T cells (Isoda et al., 2017). The *BCL11b* enhancer embedded in the *ThymoD* DNA locus repositions from the nuclear lamina to the interior prior to the activation of *BCL11b*. Artificial insertion of a polyA signal downstream of the TSS of *ThymoD* maintained the silenced local chromatin state and inhibited its repositioning into the nuclear interior, consequently leading to failed activation of *BCL11b* (Isoda et al., 2017).

It is possible that alternative enhancer usages may contribute to the sustained expression of *HAND2* in *Hand2as*^F/F^ embryos. In fact, enhancer redundancy, which is commonly observed for expression of essential developmental genes, has been suggested to provide phenotypic robustness in mammalian development (Osterwalder et al., 2018). Interestingly, HiCap analysis in mESCs (Sahlen et al., 2015) showed that the *HAND2* promoter interacts with both upstream and downstream DNA elements embedded in the *Hand2as* and *5033428I22Rik* lncRNA loci. Loss of the entire *Hand2as* locus might lead to alternative engagement of the *HAND2* promoter with downstream enhancers, which may sustain but not precisely control *HAND2* expression. A recent study of promoter competition between the lncRNA *Pvt1* and its nearby oncogene *MYC* for engagement with intragenic enhancers embedded in the *Pvt1* locus demonstrated the importance of proper enhancer-promoter interactions in regulating the precise level of *MYC* expression in breast cancer cells. Transcription interference of the *Pvt1* promoter enhances *MYC* expression and cancer cell growth *in vivo* (Cho et al., 2018).

Expression changes of genes involved in muscle contraction that are distal to the *Hand2as* locus appeared to be more pronounced than *HAND2* expression changes in bulk analysis of the whole hearts or ventricles from *Hand2as*^F/F^ mutants. These subtle *cis* effects could be easily missed in conventional ensemble analysis, thus confounding the mechanistic interpretation of a lncRNA’s function in *cis* or in *trans.* Although the molecular effects of *Hand2as* on *HAND2* expression in individual cells were moderate, these changes were prevalently and robustly detected in hundreds of cells across all four cardiac cell types, and were also observed in the septum and the right atrium of E16.5 *Hand2as*^F/F^ embryos. These results emphasize the usage of single-cell approaches to delineate the primary molecular effects of lncRNA inhibition in heterogeneous cell populations. Like many lncRNAs such as *Flicr* and *Fendrr* (Grote et al., 2013; Zemmour et al., 2017), *Hand2as* deletion had subtle molecular effects which nonetheless resulted in profound biological consequences *in vivo*.

*HAND2* expression undergoes a decrease after E10.5 and then remains low throughout the remaining course of heart development (Srivastava et al., 1997; Tamura et al., 2014). The observed upregulation of *HAND2* in E11.5 and E16.5 *Hand2as*^F/F^ embryos may reflect improper downregulation of *HAND2* during cardiac development. Interestingly, studies of various mouse models have reported that aberrantly high levels of embryonic *HAND2* led to ventricular septal defects (VSDs) (Supplementary file 1). For example, *MYH7*-driven overexpression of *HAND2* prevented the formation of the interventricular septum in embryos (Togi et al., 2006). Overdosage of *HAND2* is reportedly a major cause of perinatal lethality and heart phenotypes, including severe VSDs, in the *Rim4* mouse model which mimics a human chromosomal disorder caused by partial trisomy of distal 4q (4q+), a region containing 17 genes including *HAND2* (Tamura et al., 2013). Mice deficient in *miRNA-1-2* expressed more HAND2 protein, and exhibited a VSD and embryonic death from E15.5 to just after birth with 50% penetrance (Zhao et al., 2007). Moreover, *MYH6*-driven overexpression of *HAND2* in adult cardiac muscle cells caused pathological heart hypertrophy (Dirkx et al., 2013). Comparisons of the above mouse models with the three *Hand2as* KO mice we generated suggest a correlation between the level of *HAND2* overexpression and the severity of heart defects. Subtle yet prevalent upregulation of *HAND2* in *Hand2as*^F/F^ embryonic hearts did not affect the overall formation of an organized four-chamber heart; however, the morphological defects in specific regions of the heart, including lesions in the interventricular septum and ventricle hypoplasia, were moderate but sufficient to impair heart function, leading to immediate death of the mutant animals upon birth.

Based on these lines of evidence, we interpreted the observed molecular changes in cardiac gene programs as well as the morphological and functional heart defects to be a result of excess amounts of *HAND2* transcripts in *Hand2as*^F/F^ embryos. We propose a *cis*-functional role for the *Hand2as* locus in dampening *HAND2* expression to restrain cardiomyocyte proliferation, thereby leading to a balanced development of cardiac cell lineages. This study reinforces the notion that many lncRNAs act as local regulators to modulate the precise expression of genes of essential developmental importance, which may have profound physiological or pathological consequences in particular contexts upon lncRNA inhibition. Lastly, the fact that different knockout strategies produce distinct phenotypes underlines the requirement to utilize complementary genetic approaches to study the physiological functions of a lncRNA in mouse models (Han et al., 2018; Luo et al., 2016; Yin et al., 2015). We believe that careful genetic dissection coupled with single-cell analysis will lead to in- depth understanding of the functions and mechanisms of action of lncRNAs, thus truly impacting on our understanding of the noncoding genome in animal development, fitness and disease.

## Materials and Methods

### Animals

All mice we used were C57BL/6 background, with age described in the manuscript. Embryos were isolated at the developmental stages indicated in the manuscript. All animal experiments were conducted in accordance of institutional guidelines for animal welfare and approved by the Institutional Animal Care and Use Committee (IACUC) at Tsinghua University.

### Cells

HEK 293T cells (CRL-3216, ATCC) were cultured in DMEM supplemented with 10% heat- inactivated fetal bovine serum (Hyclone) and 1% Penicillin/Streptomycin (Cellgro). ESCs (46C, Austin Smith Lab) were grown in DMEM (Cellgro) supplemented with 15% heat-inactivated fetal bovine serum (Hyclone), 1% Glutamax (GIBCO), 1% Penicillin/Streptomycin (Cellgro), 1% nucleoside (Millipore), 0.1mM 2-mercaptoethanol (GIBCO), 1% MEM nonessential amino acids (Cellgro), and 1000U/ml recombinant LIF (leukemia inhibitory factor) (Millipore) on gelatin-coated ^plates. All cultured cells were maintained in a humidified incubator at 37°C with 5% CO_2_ (Luo et al.,^ 2016).

### CRISPR/Cas9-mediated genetic deletion

CRISPR/Cas9-meditated genetic deletion for lncRNA knockout mice generation was performed as previously described with minor modifications (Han et al., 2014a; Han et al., 2018). Briefly, Cas9 mRNA and sgRNAs were co-injected into mouse zygotes. For each genetic deletion, we used 2 sgRNAs (*Hand2as*^P^, sg1 and sg2; *Hand2as*^D^, sg3 and sg4; *Hand2as*^F^, sg1 and sg5) (Supplementary file 9). For *Hand2as*^P^, we deleted a 1-kb DNA sequence covering the core promoter and the first two exons of *Hand2as*, with 94% of *Hand2as* DNA sequences remaining intact. When genetic deletion was confirmed, the germline transmission was performed for 2 generations by mating with C57BL/6. F2 mice and later generations were used for heterozygote intercrosses.

### Genotyping

KO primers (F and R) were designed outside of the deleted region. For WT bands amplification, we used one of the KO primers together with a primer designed inside of the deleted region (Supplementary file 9). KO bands were confirmed by Sanger sequencing (Han et al., 2018).

### Echocardiography

Echocardiography was performed on *Hand2as*^D/D^, *Hand2as*^P/P^ and their littermate control mice at 6-10 weeks old. Briefly, mice were gently restrained in the investigator’s hand during echocardiography detection. Two-dimensional, short-axis views of the left ventricle were obtained for guided M-mode measurements of the left ventricular (LV) internal diameter at end diastole (LVIDd) and end systole (LVIDs). LV internal diameter were measured in at least three beats from each projection and averaged. The fractional shortening (FS) was calculated by the following formula: FS (%) = [(LVIDd − LVIDs) / LVIDd] ×100, which represents the relative change of left ventricular diameters during the cardiac cycle.

### Histology analyses

For hematoxylin-eosin staining (HE staining), E16.5 embryonic hearts were fixed in 4% PFA overnight at room temperature, dehydrated through a graded ethanol series (50%, 75%, 90%, 95%, 100%) and paraffin embedded. After section (7 μm), the tissues were deparaffinized in xylene and rehydrated through a graded ethanol series (100%, 95%, 75%), then stained with hematoxylin and eosin. Ratio of RV area was calculated as the RV chamber area divided by whole ventricle area. These data were measured in Adobe Photoshop CC2014 after selection of the image areas with myocardial color range. For immunostaining, E9.5 embryonic hearts were fixed, dehydrated, embedded as HE staining. After section (7 μm), the tissues were deparaffinized in xylene and rehydrated through a graded ethanol series (100%, 95%, 90%, 80%, 70%, 50%), then, the tissues were heated to retrieve epitope. E9.5 embryos were fixed in 4% PFA at 4 °C 2 h, dehydrated by 30% sucrose, embedded in OCT. Then frozen sections was cut at 7μm on a cryostat set at −20 to −25 °C. Immunostaining was performed with primary antibodies of TNNI3 (Abcam, ab56357), HAND2 (Abcam, ab200040). Primary antibodies were visualized by staining with Alexa-conjugated secondary antibodies: Alexa Fluor 488 donkey anti-goat (Life Technologies, #A-11055) and Alexa Fluor 555 donkey anti-rabbit (Life Technologies, #A-31572) with 200 fold diluted. All the slides were mounted in VECTASHIELD hardset antifade mounting medium (Vector Laboratories) and imaged on Zeiss Microsystems.

### RNA *in situ* hybridization

Whole mount *in situ* hybridization was carried out with digoxigenin-labelled antisense RNA probes as previously described with some modifications (Anderson et al., 2016; Wei et al., 2011). In brief, RNA probe for *HAND2* were amplified from cDNA of mouse embryonic heart and transcribed *in vitro* using T7 RNA polymerase (Roche, 10881767001) with DIG RNA labeling mix (Roche, 11277073910) (Supplementary file 9). Embryos were fixed in 4% PFA at 4 °C overnight, dehydrated through a graded methanol series (50%, 75%, 100%) and stored in 100% methanol at −20 °C. The ^embryos were bleached in a solution containing 30% H_2_O_2_: methanol 1:5 for 2 hr, then rinsed in^ methanol, rehydrated through a graded methanol series (100%, 75%, 50%), and then washed in PBS. The embryos were post-fixed 20 min in 4% PFA. After washed by PBS, embryos were transferred to the hybridization buffer (50% formamide, 5×SSC, 500 µg/ml yeast RNA, 50 µg/ml heparin and 0.1% Tween-20) and pre-hybridized 4 hr at 65 °C. Hybridizations were performed in fresh hybridization buffer containing 0.25 ng/µl digoxigenin-labelled antisense RNA probes overnight at 65 °C. Post- hybridization washes were performed at 65 °C by wash buffer 1 (50% formamide, 2× SSC), wash buffer 2 (2× SSC), wash buffer 3 (0.2× SSC), then performed by MABT (100 mM maleic acid, 150 mM NaCl and 0.1% Tween-20) at room temperature. After 1 hr blocking at room temperature in 10% sheep serum, 2% blocking reagent (Roche, 11096176001) (diluted in MABT), embryos were incubated overnight at 4 °C in blocking solution as above, with anti-DIG-AP antibody (Roche, 11093274910, 1:3,000). Then mouse embryos were washed in MABT at room temperature. After the post-antibody washes, embryos were washed in NTMT (100 mM NaCl, 100 mM Tris-HCl at pH 9.5, 50 mM MgCl2 and 0.1% Tween-20). Staining was realized in BM Purple AP Substrate (Roche, 11442074001).

### Ink injection

Chinese ink (Yidege) was injected into the left ventricles of E16.5 embryos, to visualize the organization of the arteries.

### Transfection

Plasmids were transfected into 293T or ESCs by lipofectamine 2000 (Life Technologies,#200059-61). For western blot, cells were harvested 24 hr after transfection. *HAND2* cDNA with different length of 5’UTR or without 5’UTR were cloned into *piggyBac* vector. *PiggyBac*-GFP were co-transfected with *HAND2* cDNA as control of transfection efficiency. For validation of HAND2 antibody, flag-tag was added to the N-terminal of HAND2.

### Western Blot

Cultured cells were washed by PBS and boiled in 5× SDS sample buffer for 5 min at 95°C. After SDS-PAGE and transfer, membranes were blocked in 5% milk/TBS-Tween. Primary antibody were applied 2 hr and secondary HRP-conjugated antibodies were applied for 1 hr at room temperature. Membranes were washed for 3× 10 min in TBS/Tween after each antibody incubation, and incubated with ECL substrate before exposure to X-ray film. Primary antibodies of HAND2 (Abcam, ab200040), β-TUBULIN (Abmart, M30109), FLAG (EASYBIO, BE2005-100) and GFP (CWBIO, CW0086) were used. Secondary antibodies used included goat anti-mouse IgG (CWBIO, CW0102) and goat anti-rabbit IgG (CWBIO, CW0103). Antibodies were used following the manufacture recommended concentration.

### RT-qPCR

Tissue were washed by PBS and harvested in TRIzol reagent (Life Techonologies, #15596018). Adult cardiomyocytes were isolated using type II collagenase in the Langendorff retrograde perfusion mode (O’Connell et al., 2007). Total RNA was extracted as the manufacture recommended procedure. 0.5 to 2 μg total RNA was used for reverse transcription by RevertAid First Strand cDNA Synthesis Kit (Fermentas, K1622) with random primers. Reverse transcription and quantitative PCR (RT-qPCR) were performed using iTaq Universal SYBR Green Supermix (Bio-Rad, 1725121) on a Bio-Rad CFX384 RealTime System. Error bars in RT-qPCR analysis represent standard error of mean expression relative to *GADPH* or *18S* expression, or average fold changes compared to the indicated control (Supplementary file 9).

### Bulk RNA-seq and data analysis

Adult (8-wk) cardiomyocytes of CTRL (*Hand2as*^+/+^ and *Hand2as*^D/+^) and *Hand2as*^D/D^, E11.5 hearts and E16.5 ventricles of *Hand2as*^F/+^ and *Hand2as*^F/F^ from heterozygotes intercrosses were subjected for RNAseq following polyA purification. The RNA libraries were constructed by following Illumina library preparation protocols. High-throughput sequencing was performed on Illumina HiSeq 2500 or HiSeq X TEN. All RNA-seq data were mapped to the mouse reference genome (mm9) using TopHat (version 2.0.11) (Trapnell et al., 2012). Reads were assigned to their transcribed strand (Tophat parameter “--library-type=fr-firststrand”). The gene expression level was calculated by Cufflinks (version 2.0.2) (Trapnell et al., 2012) using the refFlat database from the UCSC genome browser. For visualization, the read counts were normalized by computing the numbers of reads per million of reads sequenced (RPM). Gene set enrichment analysis (GSEA) (version 2.2.4) was performed by comparing mutant samples to control samples (Subramanian et al., 2005). We used gene sets from KEGG V6.0 (Kanehisa and Goto, 2000) for GSEA (Supplementary file 2). First, genes with FPKM >10 and |log2 (fold change)| > 0.2 (fold change = (FPKM of KO+0.1)/(FPKM of CTRL +0.1)) were selected as candidates. To exclude the inconsistently dysregulated candidates, we further filtered genes with t.test >0.1 or |log10 (t.test)* log2 (fold change)| <0.5 (dysregulation score). For 8-wk cardiomyocytes of *Hand2as*^D/D^, 114 and 186 genes are upregulated and downregulated, respectively (Supplementary file 3). For E11.5 hearts of *Hand2as*^F/F^, 169 and 101 genes are upregulated and downregulated, respectively (Supplementary file 4). For E16.5 ventricles of *Hand2as*^F/F^, 31 and 19 genes are upregulated and downregulated, respectively (Supplementary file 5). Heatmaps were drawn by Cluster 3.0 and viewed by Treeview. The colors represent the fold change of gene expression which is relative to average FPKM of each gene across all analyzed samples.

### Single-cell RNA-seq and data analysis

E11.5 hearts of wild type and *Hand2as*^F/F^ from heterozygotes intercrosses were subjected for Single-cell RNA-seq. We harvested six embryonic hearts for each genotype. Embryonic hearts were trypsinized (0.25%, 5min, 37°C) individually, and subjected for Fluorescence-activated cell sorting (FACS) after 7-Aminoactinomycin D (AAT Bioquest, 17501) staining for collection of living single cells. Next, six embryonic hearts were combined as a single sample for 10X Genomics Single Cell 3’ library construction (10X Genomics, PN-120237). The RNA libraries were constructed by following the manufacture recommended procedure to get ~5,000 cells barcoded per sample. Sequencing data analyses including sample demultiplexing, barcode processing and single-cell 3’ gene counting were done by the Cell Ranger Single-Cell Software Suite (http://software.10xgenomics.com/single-cell/overview/welcome) (Zheng et al., 2017). We got ~190 million reads for 3,469 detected *Hand2as*^F/F^ cells and ~181 million reads for 2,563 detected wild type cells, which indicate an average of 60,000 reads per cell. The cDNA insert was aligned to mouse reference genome (mm10). Cells with less than 1,000 detected genes were removed. We used RandomForest approach to discriminate a population of hemopoietic cells and excluded them from downstream analysis, and finally obtained 1,492 single cells from wild type heart and 2,108 single cells from *Hand2as*^F/F^ heart. On the median, we detected 3,492 genes per cell. For comparative analysis of WT and mutant single-cell datasets, we took the union of the top 1,000 genes with the highest dispersion (var/mean) from both datasets (“WT” object and “Mutant” object) to perform the alignment procedure in the Seurat integration procedure (Butler et al., 2018). We run a canonical correlation analysis (CCA) to identify common sources of variation between the two datasets. Then we aligned the top 7 CCA subspaces (or dimensionalities) to generate a single new dimensional reduction integrated WT and mutant datasets used for subsequent analyses such as t-distributed stochastic neighbor embedding (t-SNE) visualization. Next we used the “FindClusters” function to identify four main cardiac cell types and verified them by known marker genes (Li et al., 2016). We achieved 43% cardiomycytes, 12% epicardial cells, 31% mesenchymal cells and 14% endothelial cells (Supplementary file 6). The dominant composition of CMs in our data is consistent with two previous single-cell studies which profiled 96 cells at E11.5 and 1,165 cells at E10.5 of embryonic hearts (Dong et al., 2018; Li et al., 2016). Increases percentages of MCs and EPs at E11.5 compared to those in the E10.5 heart reported previously are in accordance with increased proliferation of cushions and epicardium (Li et al., 2016). To identify unique cluster-specific marker genes and for heatmap plotting, we used the Seurat function “FindAllMarkers” (thresh.test = 0.5, test.use = “roc”) and define a group of differentially expressed genes (DEGs) containing 1750 genes (Supplementary file 7). Next, we used the function of “FindMarkers” to identify dysregulated genes between WT and mutant hearts in each cell type, which were used for Gene Ontology analysis (Supplementary file 8). Because the median UMI we detected in most of single cells did not reach one million UMIs, we used log2 (TPM/10 + 1) rather than log2 (TPM + 1), where TPM refers to transcripts per million, to normalize the expression levels for following analysis. Correlation coefficient of cardiac cells were analyzed by 1750 DEGs used for cell types clustering. We used average gene expression of all cardiomyocytes (591 single cells) from wild type sample as a standard gene expression profile, and calculated Pearson correlation of each single cell compared to the standard. GSEA was performed by comparing all single cells of *Hand2as*^F/F^ hearts to wild type samples.

### Published data collection

Published sequencing datasets used in this paper (Figure 1A) were collected from Encyclopedia of DNA Elements (ENCODE) and Gene Expression Omnibus (GEO), including DHS-seq of E11.5 heart (ENCSR932SBO), CTCF ChIP-seq of postnatal day 0 (P0) heart (ENCSR491NUM), H3K27ac ChIP-seq of E12.5 heart (ENCSR123MLY), H3K4me3 ChIP-seq of E12.5 heart (ENCSR688ZOR), polyA RNA-seq of E12.5 heart (ENCSR150CUE), total RNA-seq of P0 heart (ENCSR035DLJ) (Yue et al., 2014), Pol II (8WG16) ChIP-seq of E12.5 heart (GSM1260035) (He et al., 2014), GATA4 ChIP-seq of E12.5 heart (GSM1260026) (He et al., 2014), HAND2 ChIP-seq of E10.5 heart (GSM1891956) (Laurent et al., 2017), NKX2-5 ChIP-seq of E12.5 heart (GSM1724109) (Ye et al., 2015), and HiCap in mESC (GSE60494) (Sahlen et al., 2015).

### Quantification and statistical analysis

Results for RT-qPCR, echocardiography and ratio of RV area are shown as mean values with error bars representing the standard error (SEM), except for *Hand2as*/*HAND2* RT-qPCR results in cardiomyocytes from CTRL (*Hand2as*^+/+, D/+^) and *Hand2as*^D/D^ (shown as median with range). Replicates are indicated in the figure legends. For each comparison between two groups, statistical analysis was performed and *p* values were calculated with an unpaired two-tailed Student’s t test by GraphPad Prism 5 software. Measurement of heart chamber area were performed by Adobe Photoshop CC2014. Imaging data analyses were done by Zen 2012. For single-cell RNA-seq analyses, scatter plots (gene expression and correlation coefficient) are shown as median and interquartile range. We used Mann Whitney test for statistical analysis of gene expression and correlation coefficient for single-cell RNA-seq results. Fisher’s test is used for significance test of gene expression frequency for single-cell RNA-seq results.

## Acknowledgements

We thank W. Pu, L. Yu, G. Ou, Y. Chen, F. Tang, Q. Xi, and Shen Laboratory members for insightful discussion and critical reading. X.S. is supported by grants from the National Natural Science Foundation of China (8141101062, 20161310854 and 31471219), the National Basic Research Program of China (2017YFA0504204), and the Center for Life Sciences (CLS) at Tsinghua University. A.H. is supported by grants from the National Basic Research Program of China (2017YFA0103402), and the National Natural Science Foundation of China (31571487 and 31771607).

## Author contributions

X.S. and A.H supervised the study. X.H., J.Z., X.F., S.A., Y.L., S.L., and Y.Y. performed the experiments. X.H., Y.L, X.F. and H.Z. performed bioinformatics analysis. X.S. wrote the manuscript with help from X.H, J.Z. and A.H.

## Data availability

Bulk RNA-seq and single-cell RNA-seq data of embryonic hearts from progenies of *Hand2as*^F/+^ or *Hand2as*^D/+^ crosses have been deposited in the Gene Expression Omnibus (GEO) under accession number GSE102935.

## Supplementary files

Supplementary file 1. Summary of cardiac *HAND2* knockout/overexpression mouse models

Supplementary file 2. Gene sets for GSEA

Supplementary file 3. Dysregulated genes in 8-wk cardiomyocytes of *Hand2as*^D/D^ mice

Supplementary file 4. Dysregulated genes in E11.5 hearts of *Hand2as*^F/F^ embryos

Supplementary file 5. Dysregulated genes in E16.5 ventricles of *Hand2as*^F/F^ embryos

Supplementary file 6. Single-cell clustering

Supplementary file 7. 1750 differentially expressed genes in four types of cardiac cells

Supplementary file 8. Dysregulated genes in four cardiac cell types in E11.5 hearts of *Hand2as*^F/F^ embryos

Supplementary file 9. Oligos (sgRNAs, genotyping primers, RT-qPCR primers, ISH probe)

---

**Figure 1-figure supplement 1.**
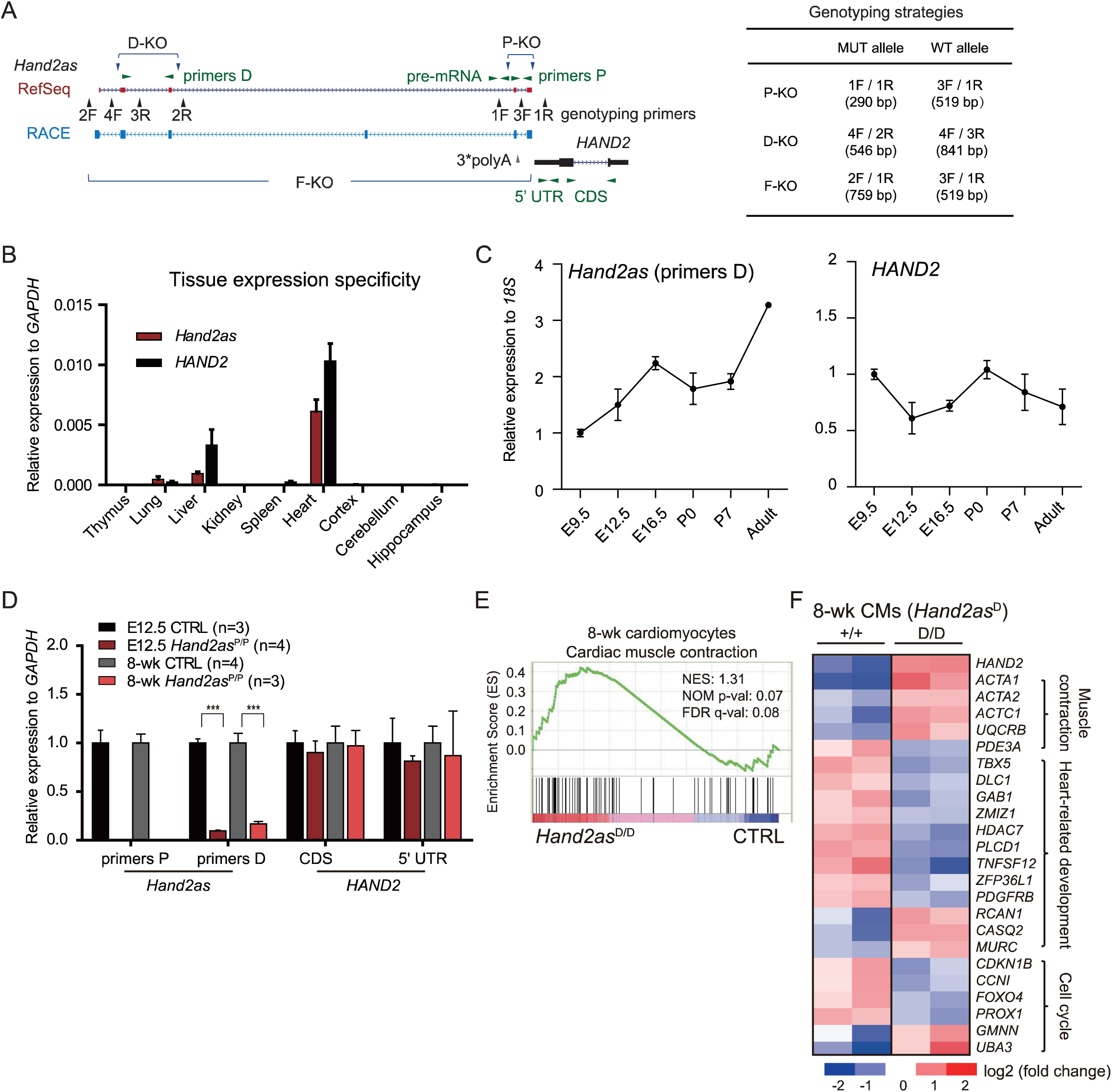
Genetic and survival characterization of *Hand2as* KO mice. (A) *Hand2as* and *HAND2* primers for RT-qPCR are shown with horizontal green arrowheads. In *Hand2as*, primers P and D are located within the proximal and distal deletions, respectively. In *HAND2*, primers 5’ UTR are located in the first 500 bp of *HAND2* mRNA, and primers CDS flank the CDS region in exon 1 and exon 2. Genotyping primers for the three *Hand2as* KO lines are shown with vertical black arrowheads. The three deletion alleles were characterized by PCR with different combinations of genotyping primers, as shown in the right panel. (B) RT-qPCR analysis of *HAND2* and *Hand2as* expression (primers D) across the indicated tissues in adult mouse (8-weeks old). The y axis shows expression relative to *GAPDH*. Data are shown as mean ± SEM. (C) RT-qPCR analysis of *HAND2* and *Hand2as* expression (primers D) during development from E9.5 to adult (8-wk). The y axis shows expression relative to *18S* (normalized to *Hand2as* or *HAND2* expression at E9.5). Data are shown as mean ± SEM. In embryos, cardiac expression of *Hand2as* continues to rise, while that of *HAND2* decreases from E9.5 to E16.5. Similarly, after birth, *Hand2as* shows significantly increased expression from postnatal day P0 to adult (8-wk), while *HAND2* expression appears to be slightly decreased. (D) RT-qPCR analysis of *Hand2as* and *HAND2* in E12.5 hearts and 8-wk cardiomyocytes from the control (CTRL, *Hand2as*^+/+, P/+^) and *Hand2as*^P/P^ mice. The y axis shows expression relative to *GAPDH* (normalized to CTRL). Data are shown as mean ± SEM. ***, p<0.001. (E) Gene Set Enrichment Analysis (GSEA) of 8-wk cardiomyocytes from CTRL (*Hand2as*^+/+, D/+^) (n=7) and *Hand2as*^D/D^ (n=6) mice. Enrichment plot shows that genes related to cardiac muscle contraction (KEGG PATHWAY: mmu04260) are upregulated in *Hand2as*^D/D^ mice compared to controls. (F) Heatmap showing representative genes that are dysregulated in 8-wk cardiomyocytes of *Hand2as*^D/D^ (rep1 and rep2) compared to wild-type (rep1 and rep2) littermates (n=2). Genes shown are involved in muscle contraction, heart-related development and cell cycle. The colored bar indicates log2 (fold change over average FPKM of each gene across 4 samples). Other samples that are not in the same litter are not considered. n, number of analyzed mice; NES, Normalized Enrichment Score.

**Figure 2-figure supplement 1.**
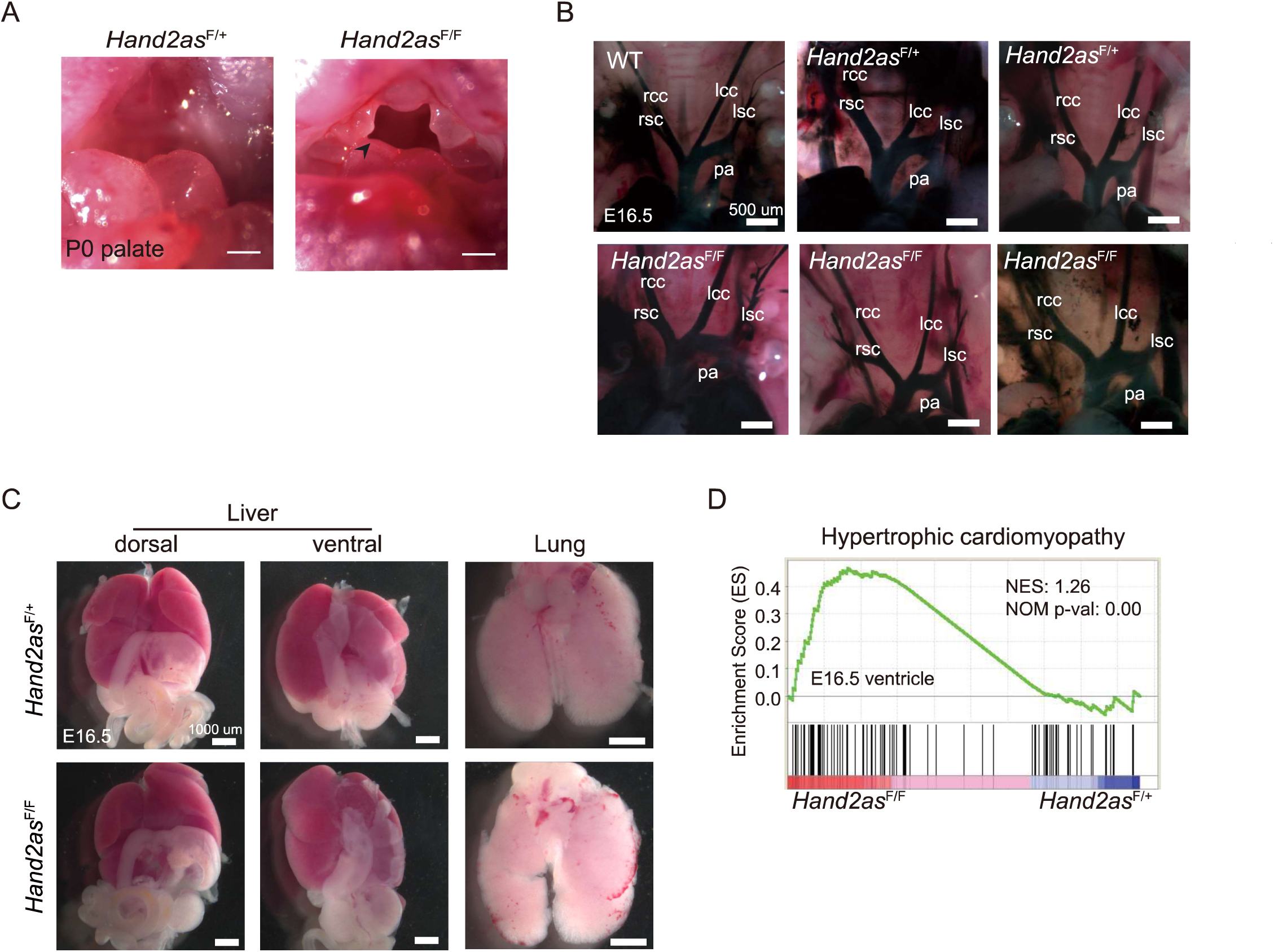
Morphological characterization of *Hand2as*^F/F^ embryos. (A) Gross morphological analysis of P0 palate of *Hand2as*^F/+^ and *Hand2as*^F/F^ mice reveals craniofacial defects (arrowhead) in *Hand2as*^F/F^ newborns. A suckling defect is not the cause of death in *Hand2as*^F/F^ pups, as these animals die immediately after birth. Mice lacking the branchial arch enhancer have craniofacial defects and cannot suckle, and die with an empty stomach 24 hours after birth. (B) Vasculature visualization by Chinese ink injection into the left ventricle of *Hand2as*^F/F^ and CTRL (*Hand2as*^+/+^ and *Hand2as*^F/+^) embryos at E16.5. lcc, left common carotid artery; lsc, left subclavian artery; pa, pulmonary artery; rcc, right common carotid artery; rsc, right subclavian artery. Scale bar, 500 µm. (C) Gross morphological examination of liver and lung of *Hand2as*^F/+^ and *Hand2as*^F/F^ embryos at E16.5. Scale bar, 1000 µm. (D) Gene Set Enrichment Analysis (GSEA) of E16.5 ventricles of *Hand2as*^F/+^ and *Hand2as*^F/F^ mice (n=3 for each genotype). Enrichment plot shows that genes related to hypertrophic cardiomyopathy (KEGG PATHWAY: mmu05410) are upregulated in *Hand2as*^F/F^ mice compared to *Hand2as*^F/+^ controls.

**Figure 3-figure supplement 1.**
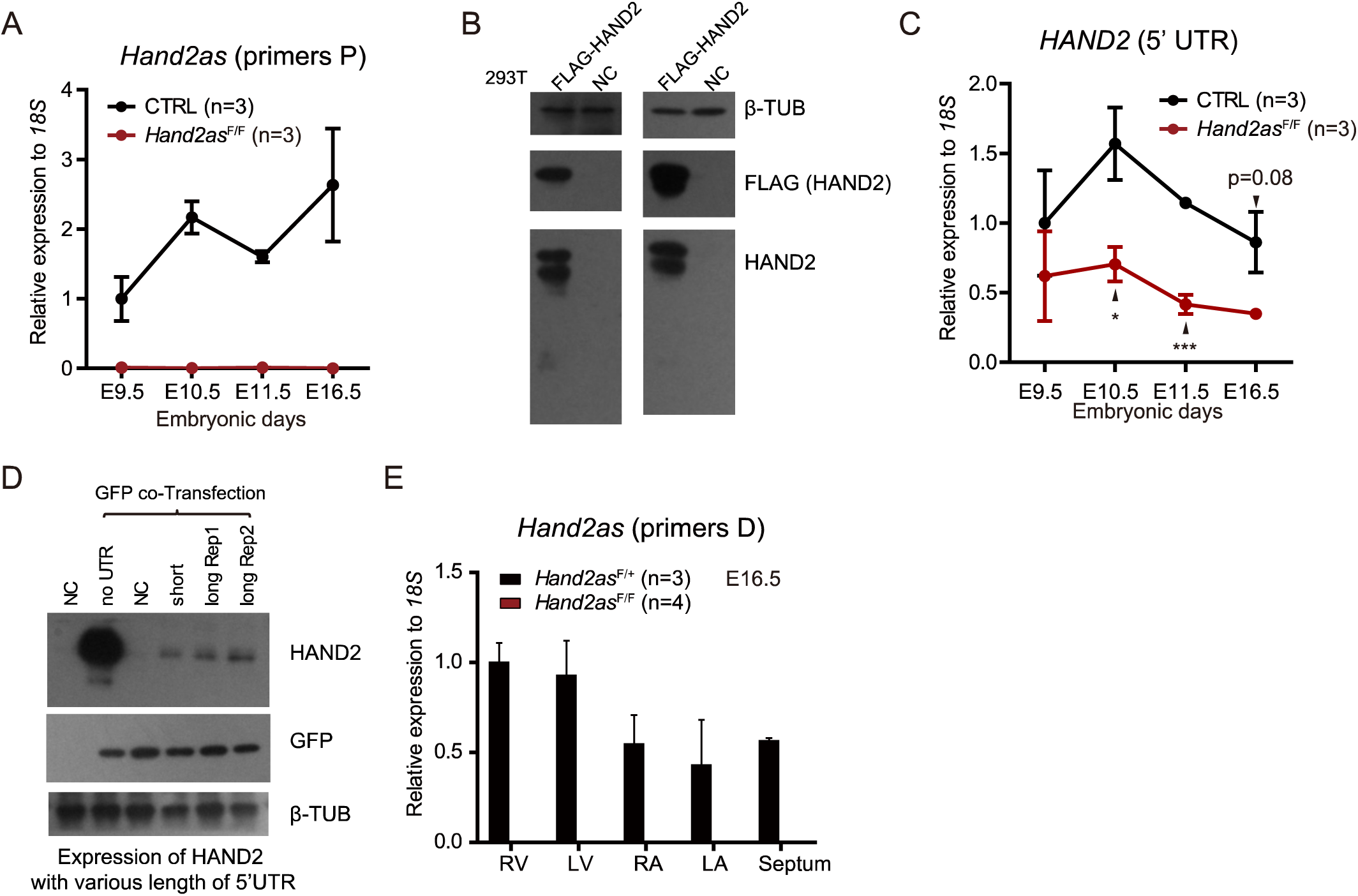
Molecular analysis of *Hand2as*^F/F^ embryos. (A) RT-qPCR analysis of *Hand2as* (primers P) in embryonic hearts at various stages from CTRL (*Hand2as*^+/+, F/+^) and *Hand2as*^F/F^. The y axis shows expression relative to *18S* (normalized to *Hand2as* expression in CTRL embryos at E9.5). Data are shown as mean ± SEM. (B) Western blot validation of the HAND2 antibody in 293T cells transfected with Flag-HAND2. β-TUBULIN is the loading control. (C) RT-qPCR analysis of *HAND2* (5’ UTR) in embryonic hearts at various stages from CTRL (*Hand2as*^+/+, F/+^) and *Hand2as*^F/F^. The y axis shows expression relative to *18S* (normalized to *HAND2* 5’ UTR expression in CTRL embryos at E9.5). Data are shown as mean ± SEM. *, p<0.05; ***, p<0.001. (D) Western blot shows HAND2 expression from constructs with various lengths of *HAND2* 5’UTR or without the 5’UTR. Transfections were performed in mouse ESCs. β-TUBULIN is the loading control and GFP is the transfection control. (E) RT-qPCR analysis of *Hand2as* (primers D) in dissected E16.5 embryonic hearts (RV, LV, RA, LA and septum) from *Hand2as*^F/+^ and *Hand2as*^F/F^. The y axis shows expression relative to *18S* (normalized to *Hand2as* expression in RV of *Hand2as*^F/+^ embryos). Data are shown as mean ± SEM. n, number of analyzed mice.

**Figure 5-figure supplement 1.**
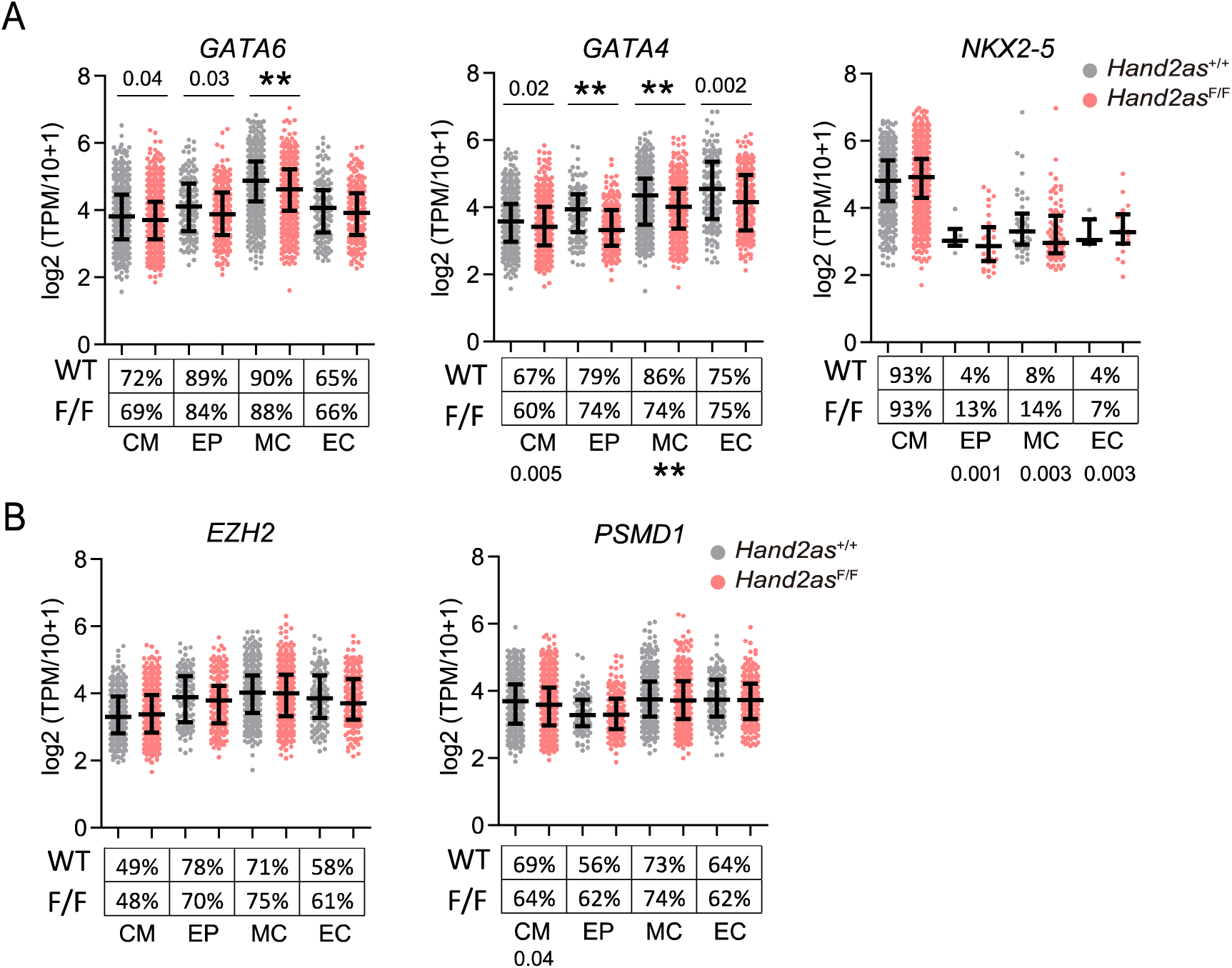
Gene expression in *Hand2as*^F/F^ embryos. (**A-B**) Scatter plots show expression level and frequency of *GATA6*, *GATA4* and *NKX2-5* (**A**), and *EZH2* and *PSMD1* (**B**) in cardiac cells of *Hand2as*^F/F^ and wild-type embryonic hearts. Data are shown as median and interquartile range. The *p*-values are indicated: 0.0001<*p*<0.05; **, *p*<0.0001. The percentages of expressing cells (TPM>0) in each cell type are shown under the plots.

**Figure 6-figure supplement 1.**
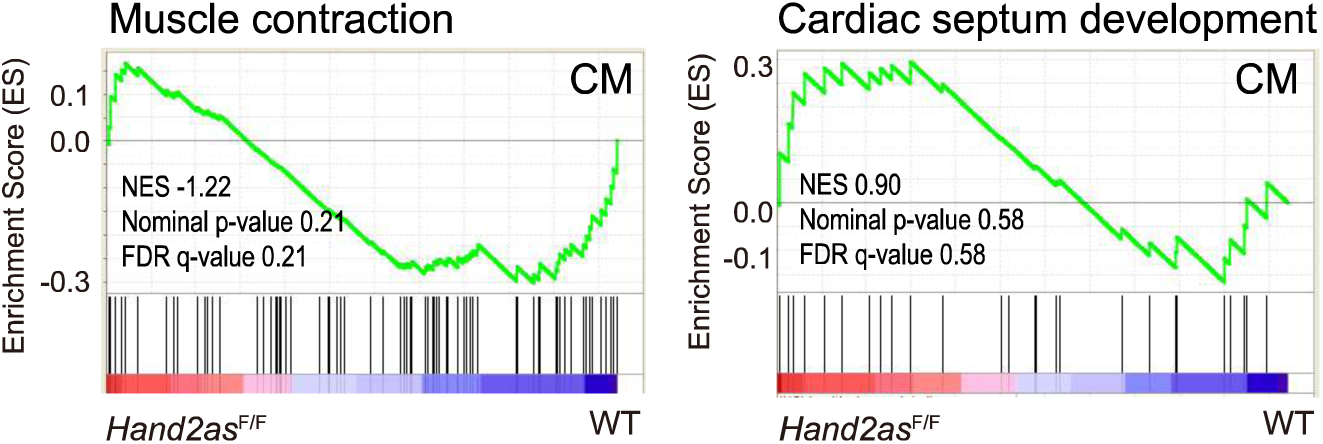
Aberrant cardiac gene programs in *Hand2as*^F/F^ embryos. Non-significant changes of genes related to muscle contraction (GO: 0006936) and cardiac septum development (GO: 0003279) in CMs of *Hand2as*^F/F^ versus wild-type E11.5 hearts by GSEA.

